# Recapitulating bone development for tissue regeneration through engineered mesenchymal condensations and mechanical cues

**DOI:** 10.1101/157362

**Authors:** Anna M. McDermott, Samuel Herberg, Devon E. Mason, Hope B. Pearson, James H. Dawahare, Joseph M. Collins, Rui Tang, Amit N. Patwa, Mark W. Grinstaff, Daniel J. Kelly, Eben Alsberg, Joel D. Boerckel

**Affiliations:** McKay Orthopaedic Research Laboratory, Department of Orthopaedic Surgery, Perelman School of Medicine, University of Pennsylvania, Philadelphia, PA.; Department of Aerospace and Mechanical Engineering, University of Notre Dame, Notre Dame, IN.; Department of Mechanical Engineering, Trinity Center for Bioengineering, Trinity College Dublin, Dublin, Ireland.; Department of Biomedical Engineering, Case Western Reserve University, Cleveland, OH.; Wake Forest Institute for Regenerative Medicine, Wake Forest School of Medicine, Winston-Salem, NC, USA.; Department of Bioengineering, University of Pennsylvania, Philadelphia, PA.; Department of Biomedical Engineering, Boston University, Boston, MA.; Department of Orthopaedic Surgery, Case Western Reserve University, Cleveland, OH.; National Center for Regenerative Medicine, Division of General Medical Sciences, Case Western Reserve University, Cleveland, OH.; Current address: Departments of Ophthalmology | Cell and Developmental Biology Biochemistry and Molecular Biology, SUNY Upstate Medical University, Syracuse, NY.

## Abstract

Large bone defects cannot heal without intervention and have high complication rates even with the best treatments available. In contrast, bone fractures naturally healing with high success rates by recapitulating the process of bone development through endochondral ossification.^1^ Endochondral tissue engineering may represent a promising paradigm, but large bone defects are unable to naturally form a callus. We engineered mesenchymal condensations featuring local morphogen presentation (TGF-β1) to mimic the cellular organization and lineage progression of the early limb bud. As mechanical forces are 2,3 critical for proper endochondral ossification during bone morphogenesis^2,3^ and fracture healing, we hypothesized that mechanical cues would be important for endochondral regeneration.^4,5^ Here, using fixation plates that modulate ambulatory load transfer through dynamic tuning of axial compliance, we found that *in vivo* mechanical loading was necessary to restore bone function to large bone defects through endochondral ossification. Endochondral regeneration produced zonal cartilage and primary spongiosa mimetic of the native growth plate. Live human chondrocytes contributed to endochondral regeneration *in vivo*, while cell devitalization prior to condensation transplantation abrogated bone formation. Mechanical loading induced regeneration comparable to high-dose BMP-2 delivery, but without heterotopic bone formation and with order-of-magnitude greater mechanosensitivity.^6–8^ *In vitro*, mechanical loading promoted chondrogenesis, and upregulated pericellular collagen 6 deposition and angiogenic gene expression. Consistently, *in vivo* mechanical loading regulated cartilage formation and neovascular invasion dependent on load timing. Together, this study represents the first demonstration of the effects of mechanical loading on transplanted cell-mediated bone defect regeneration, and provides a new template for recapitulating developmental programs for tissue engineering.

---

Long bone morphogenesis is initiated by condensation of mesenchymal cells in the early limb bud, which differentiate and mature into the cartilaginous anlage that gives rise to endochondral bone formation. This process is dependent on both local morphogen gradients and mechanical forces *in utero*.^3,9^ Natural bone fracture healing recapitulates endochondral bone development, but only under conditions of compressive interfragmentary strain.^5,10^ Without mechanical loading, fractures will heal through direct, intramembranous bone formation,^11^ implicating mechanical cues as critical regulators of endochondral ossification. The emerging paradigm of biomimetic bone tissue engineering aims to replicate the endochondral process,^12,13^ but functional endochondral bone regeneration using transplanted human progenitor cells remains elusive potentially due to insufficient recapitulation of the essential cellular, biochemical, and mechanical stimuli. Here, we engineered scaffold-free human bone marrow-derived mesenchymal stem cell (hMSC) condensations through cellular self-assembly into sheets^14^. Morphogen-releasing gelatin microspheres were incorporated into the condensations to sustained the local presentation of transforming growth factor-β1 (TGF-β1)^14^ and induce endochondral lineage progression upon implantation. We controlled *in vivo* mechanical loading through dynamic modulation of fixation plate compliance.^6,15^

Engineered mesenchymal condensations were self-assembled into sheets of 2 × 10^6^ cells containing local presentation of 600 ng rhTGF-β1. After two days culture, the condensations exhibited homogeneous cellular organization without histologically detectable sulfated glycosaminoglycan (sGAG) deposition or bone formation (Fig. 1a, Extended Data Fig. 1a). Cellular organization was qualitatively similar to that of the developing mouse limb bud at E11.5-12.5 (Fig. 1b, Extended Data Fig. 1b). Local TGF-β1 presentation upregulated and sustained expression of Sox9, Aggrecan, and Collagen 2a1 (Fig. 1c) at the message-level, with minimal expression of osteogenic markers (Osterix, Runx2, Alkaline phosphatase, and Collagen 1a1). TGF-β1 also increased protein-level phosphorylation of the chondrogenic transcription factor SMAD3 at day 2 *in vitro* (Fig. 1e,f). These data demonstrate chondrogenic lineage priming consistent with the known dynamics of TGF-β1-SMAD signaling and downstream gene expression in the developing limb at E11.5-12.5 (cf. refs. ^16–18^). After 23 days *in vitro* culture, local TGF-β1-containing condensations exhibited characteristically shaped chondrocytes and substantial sGAG matrix (Fig. 1d), demonstrating chondrogenic lineage commitment without additional growth factor supplementation in the media.

**Figure 1:**
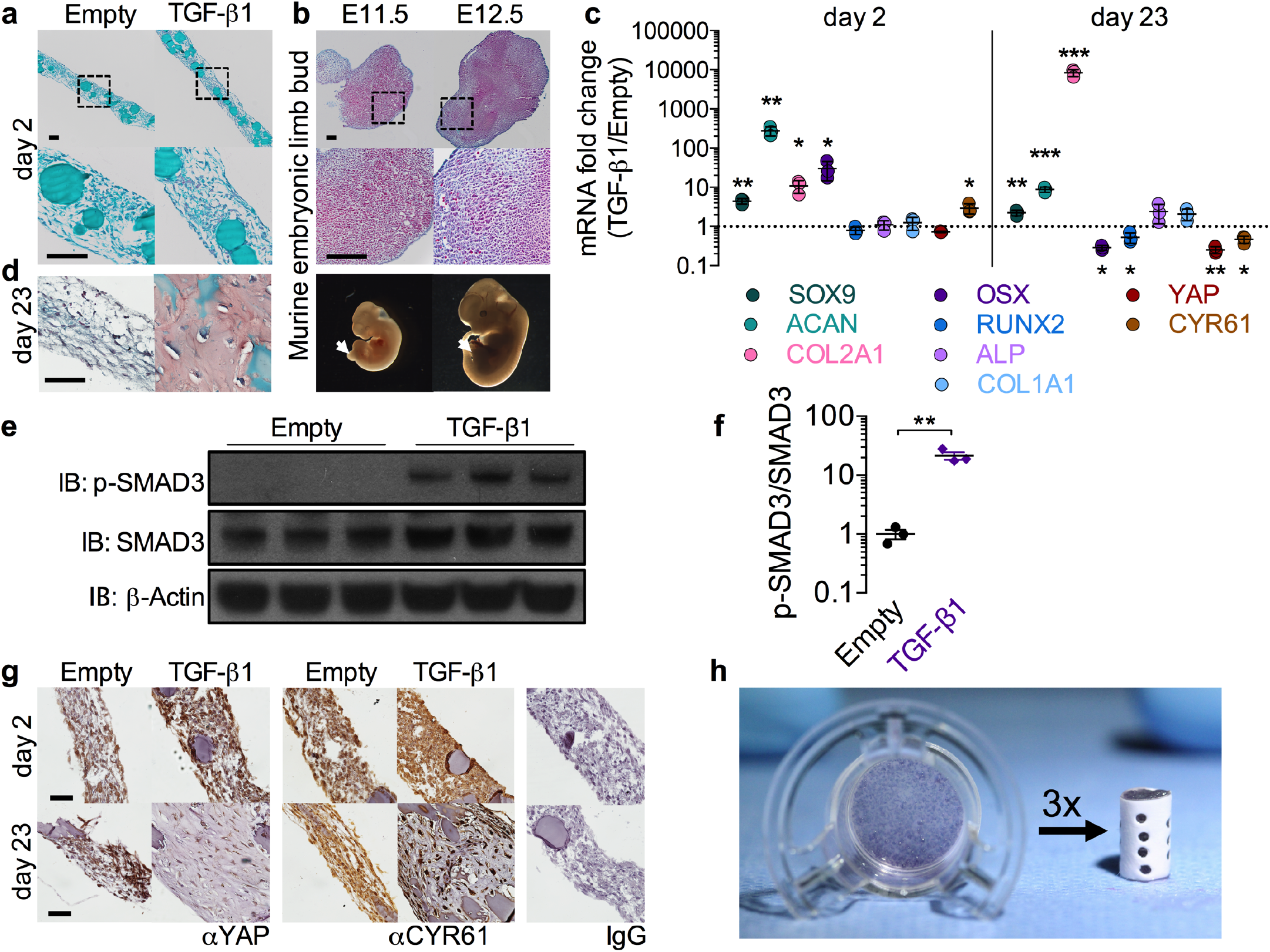
Engineered mesenchymal condensations mimic the embryonic limb bud. **a**, Safranin-O/Fast green staining of hMSC sheets with empty gelatin microspheres (left), or loaded with 600 ng rhTGF-β1 (right) cultured for two days on transwell inserts. Bottom: magnification of enclosed areas. **b**, Safranin-O/Fast green staining of murine limb buds at embryonic days 11.5 and 12.5 (E11.5, E12.5). Bottom: photomicrographs of embryos with limb buds indicated by arrows. **c**, Messenger RNA expression of chondrogenic, osteogenic, and YAP pathway genes at day 2 and day 23 *in vitro*; qRT-PCR results were normalized to GAPDH and expressed as fold-change over empty microsphere control sheets (n = 3 sheets per group). **d**, Safranin-O/Fast green staining of sheets at day 23 *in vitro*. **e**, Western blot of phosphorylated-SMAD3 activity at day 2 *in vitro* with β-actin control and **f**, band intensity of p-SMAD3/SMAD3 ratio expressed as fold change over sheets without growth factor **g**, Immunostaining for YAP and target gene CYR61 at days 2 and 23 *in vitro*. Right: negative control isotype IgG (rabbit, top; mouse, bottom). **h**, Photograph of hMSC sheet cultured 2 days on transwell insert (left) and engineered mesenchymal condensation for implantation (right), assembled from three sheets within a perforated electrospun mesh tube. *p<0.05, **p<0.01, ***p<0.001 TGF-β1 treated vs. empty, unpaired two-tailed Student’s t-test for each independent gene, with corrections for multiple comparisons by the Bonferroni method. Data shown with mean ± s.d. Scale bars, 100 μm. Histological images are representative of three independent samples per group.

While the molecular mechanisms that control endochondral ossification remain incompletely understood, recent evidence from our laboratory and others implicates the transcriptional co-activator Yes-associated protein (YAP) as a mechanosensitive,^19^ TGF-β1-inducible^20^ regulator of progenitor cell lineage specification, promoting endochondral bone development^21^ but inhibiting chondrogenesis.^22^ Consistent with these reports, we observe that TGF-β1 presentation transiently upregulated transcription of the YAP target gene cysteine-rich angiogenic inducer 61 (Cyr61) (Fig. 1c) and increased YAP protein levels at day 2 (Fig. 1g, top row), while Cyr61 expression was reduced in differentiated chondrocytes at day 23 (Fig. 1c,g, bottom row), consistent with an inhibitory role of YAP in the mature cartilage anlage.^22^ Together, these data demonstrate our capacity to mimic the early limb bud in engineered mesenchymal condensations and suggest that YAP could play a role in mechanical regulation of endochondral lineage progression during regeneration.

To evaluate the capacity of engineered mesenchymal condensations to induce endochondral bone regeneration, we created cylindrical condensations (8 mm length, 5 mm diameter) for *in vivo* implantation. These were compiled by placing three hMSC sheets (after 2 days maturation *in vitro)* into perforated electrospun polycaprolactone nanofiber mesh tubes^23^ (Fig. 1h), for a total of 6 × 10^6^ cells and 1.8 μg of rhTGF-β1 per construct. The nanofiber mesh tubes served to maintain condensation shape and were perforated to facilitate vascular invasion.^6,24^ Condensations were then implanted in critical-sized (8 mm) bone defects, surgically created in femora of athymic Rowett nude rats (RNU), as described previously.^6,25^ This 8 mm segmental defect model represents a challenging test-bed for regenerative strategies, being >60% larger than the minimum gap size necessary to prevent spontaneous repair (5 mm).^25,26^

Mechanical stimuli promote proper endochondral ossification in both bone development and fracture healing,^2,3,27^ but the effects of *in vivo* mechanical loading on transplanted cell-mediated bone repair has not been studied. Here, we modulated ambulatory load transfer *in vivo* using custom internal fixation plates capable of temporal control of axial compliance by elective unlocking (Fig. 2a).^6,7,28^ The timing and magnitude of mechanical forces imparted to the defects were controlled in three groups: stiff (control, n = 11), early (compliant plates unlocked at implantation to allow immediate loading, n = 9), and delayed (compliant plates unlocked at week 4 to initiate loading, n = 9) (Fig. 2b, Extended Data Video 1). The multi-modal mechanical behavior of the plates was assessed by *ex vivo* mechanical testing (stiff: k_axial_ = 260 ± 28 N/mm, locked compliant: k_axial_ = 250 ± 35 N/mm, unlocked compliant: k_axial_ = 8.0 ± 3.5 N/mm; mean ± s.d.; Extended Data Fig. 2). Published data on femoral loading during the rat gait cycle^29^ and rule-of-mixtures theory were used to quantify load-sharing between the fixation plates and the defect tissue. These calculations indicated that interfragmentary strains at day 0 reached 2-3% in the stiff and delayed groups, and up to 10-15% in the early group. A recent *in vivo* strain sensor study using a modified version of the stiff plates described here confirmed these numbers within one percent for the stiff group.^30^ The amount of strain induced over time is a function of the load sharing, and therefore dependent on the amount, composition, and kinetics of tissue ingrowth; however, accounting for load sharing by ingrowing bone, we estimated strains of 5-10% upon plate unlocking at week 4 in the delayed group, with all groups converging on 0.5-3% by week 12.

**Figure 2:**
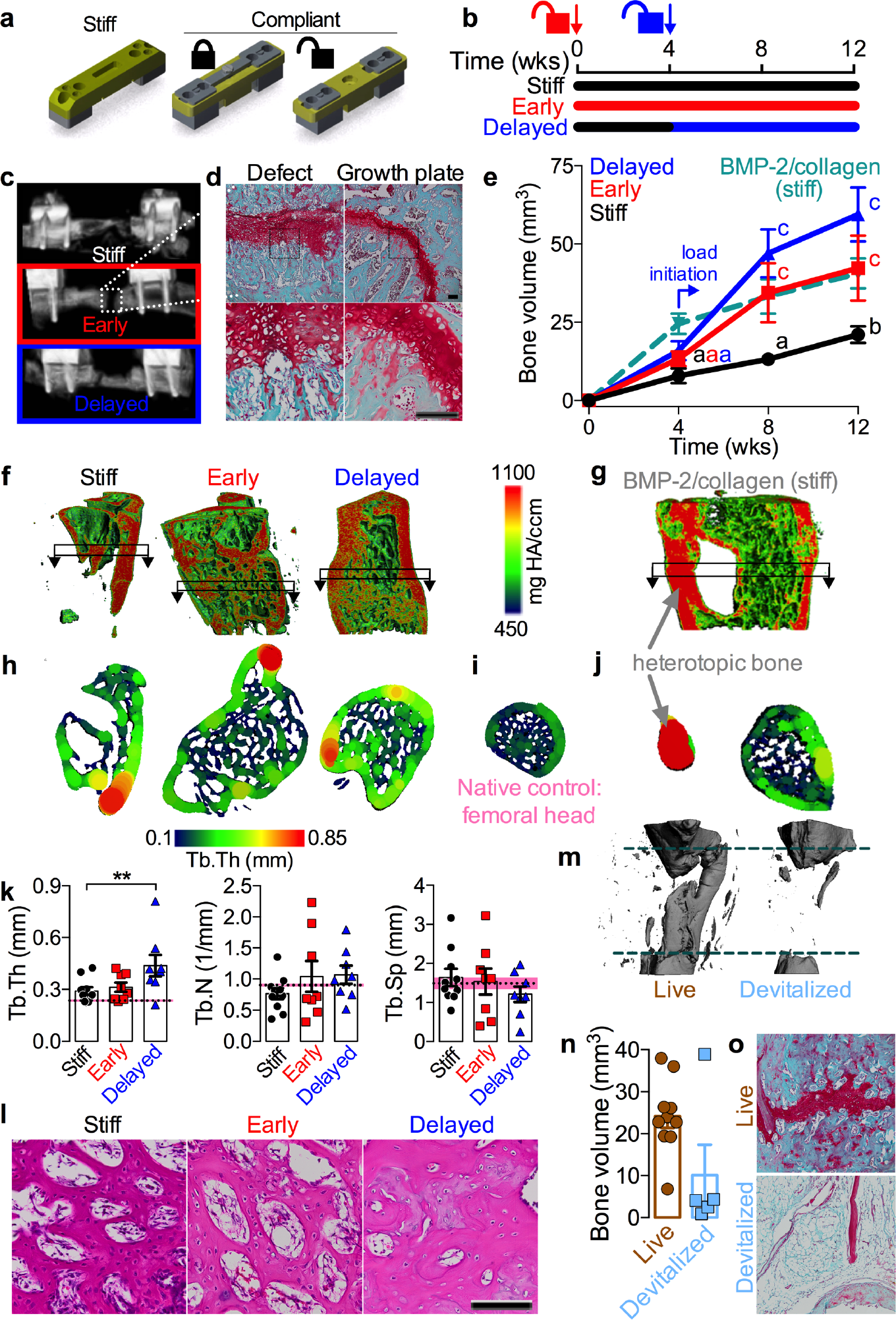
Mechanical loading enhances endochondral bone formation. **a**, Fixation plate configurations for dynamic control of ambulatory load transfer. **b**, Schematic of loading timeline with early loading featuring compliant plate unlocking at implantation and delayed loading featuring unlocking at week 4. **c**, Representative *in vivo* microCT reconstructions at week 4. Images selected based on mean bone volume at week 12. **d**, Safranin-O/Fast green staining of sagittal histological sections at week 4 (left) in comparison to the native proximal femur growth plate (right). Bottom: magnification of enclosed areas. **e**, Longitudinal microCT quantification of bone volume at week 4 (n = 11, 11, 9, and 8 for stiff, early, and delayed, and BMP-2/collagen (stiff), respectively), week 8 (n = 10, 9, 8, 8) and 12 (n = 10, 8, 8, 8). The BMP-2/collagen group featured 5μg rhBMP-2 delivered on absorbable collagen sponge. Repeated significance indicator letters (a,b,c) signify p < 0.05, while groups with distinct indicators signify p < 0.05 between groups at each time point. **f-g**, Representative 3D microCT reconstructions at week 12 with local density mapping on a virtually-cut sagittal plane. **h-i**, Local trabecular thickness mapping on transverse sections, indicated by boxed arrows in (f), in comparison to the native bone of the ipsilateral femoral head. **j**, Trabecular thickness mapping in BMP-2/collagen samples illustrating substantial heterotopic bone. **k**, MicroCT quantification of trabecular thickness (Tb.Th), number (Tb.N) and spacing (Tb.Sp) in reference to the trabecular morphology of the ipsilateral femoral head (femoral head mean ± s.d. shown as dotted line and shaded pink region). **l**, H&E-stained histological sections at week 4 (n = 1 representative sample per group). **m-o**, Effect of repeated freeze-thaw-induced construct devitalization on bone regeneration: **m**, Representative microCT reconstructions showing the original defect region bordered by dotted dark green lines. **n**, Bone volume at week 12 in live and devitalized groups (n=10, 5). **o**, Representative Safranin-O/Fast green staining at week 12 (n = 1-3 per group). All scale bars, 100 μm. All representative samples were selected to best match the average microCT morphometry of that group at the corresponding time point. Data shown with mean ± s.e.m. *p<0.05, ** p<0.01, **** p<0.0001, NS = not significant, one or two-way ANOVA with Tukey’s *post-hoc* analysis.

Bone regeneration progressed through endochondral ossification, exhibiting zonal cartilage and woven bone mimetic of the native growth plate by week 4 (Fig. 2c,d), similar to fracture callus recapitulation of development.^1^ Both early and delayed loading significantly enhanced bone formation (Fig. 2, Extended Data Figs. 3,6, Video 1) as measured by bone volume (Fig. 2e) and bone volume fraction (Extended Data Fig. 3a). Notably, loading elevated bone accumulation rate between weeks 4 and 8 (cf. slopes of bone volume vs. time curves in Fig. 2e, Extended Data Fig. 3b). This coincided with the timing of load initiation in the delayed group and the tissue differentiation stage, namely chondrocyte hypertrophy and endochondral transition in all three groups (Extended Data Fig. 4). These findings suggest that this stage of endochondral ossification is particularly responsive to mechanical forces, consistent with developmental studies showing that forces exerted by muscle contractions *in utero* are critical for proper chondrocyte hypertrophy and bone morphogenesis.^2,3^ Though early loading significantly enhanced bone formation (Fig. 2e), the bone volume fraction response in this group was highly variable (Extended Data Fig. 3a-c) and exhibited a significantly lower bridging rate compared to delayed loading (2/8 vs. 6/8, p<0.05 Chi-square test, Extended Data Fig. 3d,e) due to persistent regions of nonmineralized cartilage and fibrocartilage (Fig. 2c, Extended Data Fig. 4), similar to the pseudarthroses induced by large-deformation cyclic bending.^31^ In contrast, delayed loading induced robust bone formation (Fig. 2e), with consistent bridging rate (Extended Data Fig. 3d,e).

As a clinically-relevant positive control, a healing dose of rhBMP-2 (5 μg/defect),^32^ delivered on absorbable collagen sponge with stiff fixation, was also evaluated (n=8) (“BMP-2/collagen,” dashed line in Fig. 2e). This treatment produced inverted bone formation kinetics to the endochondral bone, with rapid bone accumulation until week 4, and reduced bone formation rate thereafter (Fig. 2e). BMP-2-treated defects did not exhibit cartilage formation (Extended Data Fig. 5a,b), but as reported clinically, BMP-2 treatment induced extensive heterotopic bone formation (Fig. 2g,j). Surgeries for the BMP-2-treated samples were performed at a separate time and therefore were not compared statistically with the other groups.

Notably, the response to mechanical loading observed for endochondral regeneration (181% increase in bone volume compared to control) was an order-of-magnitude greater than that reported previously by our group^6,7^ and others^8^ for *in vivo* loading of BMP-2-mediated repair (18-20% increase). Quantitative densitometry (Fig. 2f, Extended Data Fig. 6d) and region of interest analyses (Extended Data Fig. 6a,b) revealed mineral concentration at the defect periphery, indicative of a cortical shell in all three groups. The bone formed within this cortex exhibited well-defined trabecular architecture, which was quantitatively similar to native femoral head trabecular bone as assessed by microCT morphometry (Fig. 2h,i,k, Extended Data Fig. 6c) and histology (Fig. 2l, Extended Data Fig. 4).

Transplanted MSCs commonly exhibit rapid cell death due to lack of vascular and nutrient supply;^25^ however, in studies using an endochondral paradigm, viable donor cells have been observed up to several weeks after implantation.^33–36^ Further, recent reports suggest that hypertrophic chondrocytes may also transdifferentiate into bone-forming cells.^37–39^ Therefore, to test whether the transplanted cells functionally contributed to bone repair, we prepared identical mesenchymal condensations (6 × 10^6^ cells and 1.8 μg of rhTGF-β1 per construct) for implantation after devitalization by freeze-thaw cycling. Using stiff plates, these elicited substantially reduced bone formation compared to live cell controls (Fig. 2m-o, Extended Data Video 1) with fibrotic tissue filling the defect (Fig. 2o). Comparisons with the devitalized samples were not assessed statistically due to surgical operation at a separate time, but suggest a functional role of the transplanted cells.

As the principal test of any engineered tissue needs to be its functionality,^40^ we evaluated the degree of restoration of bone mechanical properties by torsion testing to failure at week 12, in comparison with age-matched intact femurs (Fig. 3). Despite enhanced bone formation, early loading failed to restore mechanical properties, while delayed loading significantly increased torsional stiffness and maximum torque at failure compared to stiff plate controls (Fig. 3a,b) and restored the torsional stiffness to that of intact limbs (dotted line/gray shading indicate mean ± s.d.). The corresponding structural properties, average polar moment of inertia (pMOI) and minimum pMOI (Fig. 3c,d) were also significantly elevated by delayed loading. To identify the most likely combination of factors that predict the mechanical behaviour, we performed a best-subsets analysis using average and mimimum pMOI, bone volume, binary bridging score (indicated in Fig. 3 by shaded vs. open data points; Extended Data 3d,e), and average mineral density as parameters. The most optimal model was determined by minimization of the Akaike’s information criterion (AIC).^41^ We then performed Type II multivariate regression analyses to determine the amount of variability in the measured stiffness and maximum torque that is explained by the selected predictors (R^2^).

**Figure 3:**
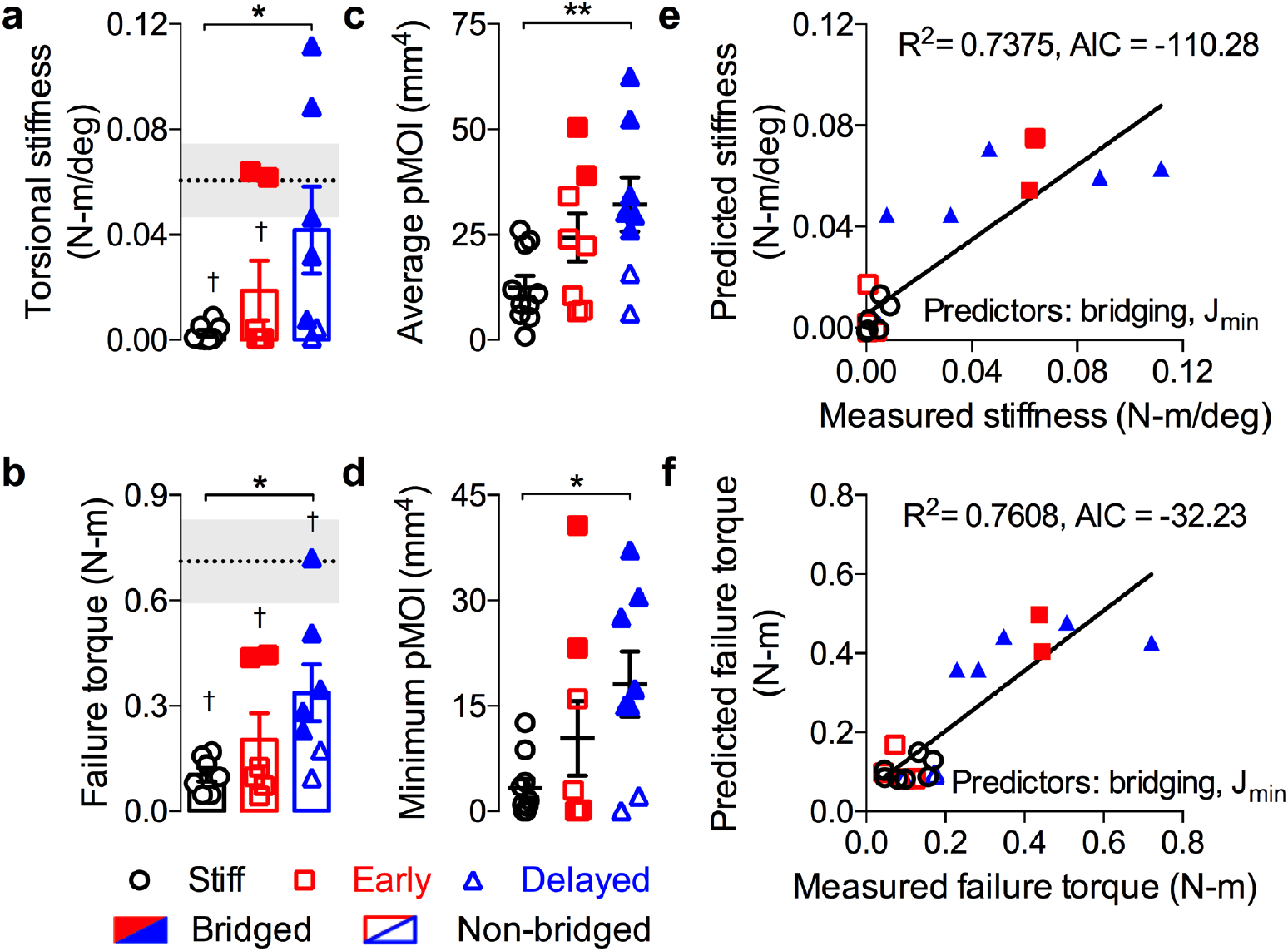
Restoration of mechanical function. Structural mechanical properties were measured by torsion to failure following microCT analysis of cross-sectional bone distribution at 12 weeks. Age-matched intact bone properties are shown as dotted line/gray shading indicating mean ± s.e.m. Samples with full defect bridging are shown in filled data points, while open data points indicate non-bridged samples. **a**, Analysis of torsional stiffness **b**, maximum torque at failure **c**, average polar moment of inertia (pMOI) and **d**, minimum pMOI (n = 8, 7, 7 for stiff, early, and delayed, respectively). Best subsets regression analysis with lowest AIC value for measured and predicted torsional stiffness (**e**) and maximum torque at failure (**f**) indicating significant contributions of minimum pMOI (J_min_) and binary bridging score. Bar graphs show mean ± s.e.m. with individual data points. Statistical comparisons between groups for each measure were performed by one-way ANOVA with Tukey’s *post-hoc* analyses, *p<0.05; †p<0.05 vs. intact bone.

For both maximum torque and torsional stiffness, minimum pMOI and bridging score were the best combined predictors from among the top 5 models (Fig. 3d,e, Extended data fig. 7). This analysis reveals that the mechanical properties were determined both by whether the defects were fully bridged with bone and the amount of the bone at the location of failure. It further explains differences in mechanical outcomes between the early and delayed loading groups; while both loading conditions significantly enhanced bone volume, the low bridging rate of the early group impaired functional repair. Together, these data demonstrate that mechanical cues are critical for restoration of bone form and function by endochondral ossification of engineered mesenchymal condensations.

To determine whether loading regulated endochondral lineage progression and matrix organization,^7,31^ we performed histological staining at weeks 3, 4, 7, and 12 (Fig. 4a,b, Extended Data Figs. 4, 9). The endochondral regenerate contained distinct bands of Safranin-O-positive cartilage featuring mature and hypertrophic chondrocytes producing woven bone. Extensive sGAG staining was observed at early time points (3 and 4 weeks), and calcified cartilage and bone at weeks 7 and 12 (Fig. 4a, Extended Data Figs. 4, 9). Both early and delayed loading enhanced and prolonged the chondral phase of endochondral ossification, as indicated by Safranin-O staining intensity (Fig. 4a, Extended Data Figs. 4, 9). Polarized light analysis of picrosirius red-stained sections^7^ revealed no differences in collagen organization during the initial healing period (4 wks; Extended Data Fig. 8), but both early and delayed loading decreased collagen fiber birefringence compared to the stiff controls at week 12 (Fig. 4b), suggesting an increased proportion of immature woven bone in the loaded groups through either increased new woven bone deposition or altered remodeling, similar to our prior observations with BMP-2-mediated, load-induced bone regeneration.^6,7^

**Figure 4:**
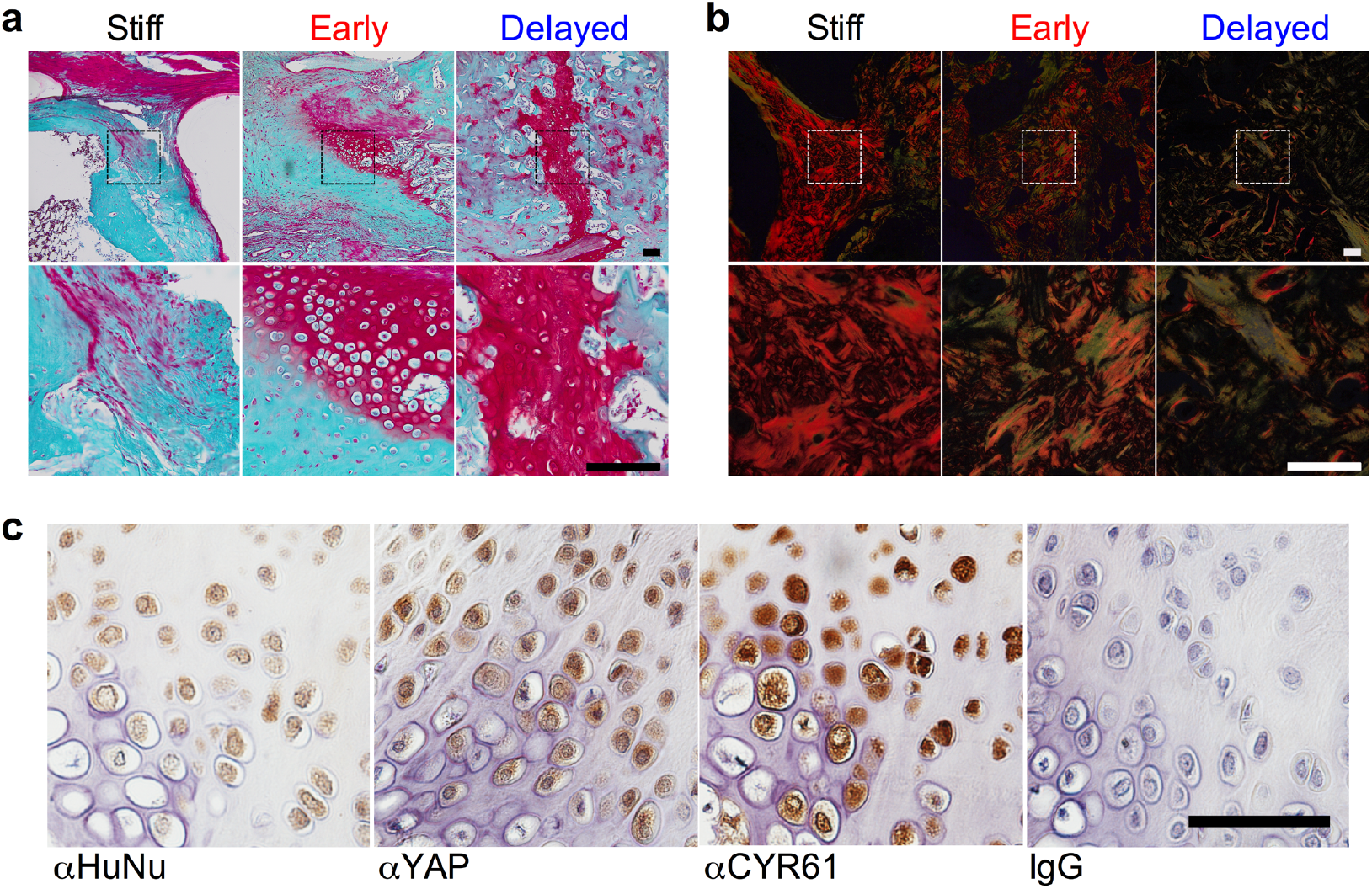
Endochondral ossification, matrix remodeling, and transplanted cell engraftment. **a**, Safranin-O/Fast green staining of sagittal histological sections of defects at week 12 (n = 1 representative sample per group). **b**, Polarized light microscopy of picrosirius red-stained histological sections at week 12. **c**, Immunohistochemistry of live-cell treated, delayed loaded defects at week 12 for human nuclear antigen (HuNu), YAP and YAP target gene CYR61 in comparison to isotype IgG negative control. Scale bar, 100 μm. All representative samples were selected to best match the average microCT morphometry of that group.

Above, we noted a functional effect of transplanted cell devitalization on regeneration (cf. Fig. 2m-o). To further elucidate transplanted cell fate and function, live human cells were immunolocalized by human nuclear antigen (HuNu) staining.^25^ Viable human cells, morphologically identifiable as mature and hypertrophic chondrocytes, actively engaged in endochondral ossification as late as week 12 *in vivo* (Fig. 4c). These human cells also exhibited nuclear-localized YAP protein and expression of the downstream angiogenic matricellular growth factor, CYR61 (Fig. 4c). Together, this suggests that the transplanted human cells both participated in endochondral lineage progression and exhibited mechanosensitive gene activity.

To determine the temporal roles of mechanical loading on chondrogenic lineage commitment, gene expression, and matrix production, we applied 10% dynamic compression *in vitro* to hMSCs encapsulated in fibrin hydrogels. Loading was applied either continuously for 5 weeks (early), for 2 weeks following a 3 week free swelling period (delayed), or for 2 weeks prior to a 3 week free swelling period (reversed), compared to 5 week free swelling (FS) controls (Fig. 5a,b). Loading significantly increased DNA content, indicating either increased proliferation or maintenance of viability (Fig. 5c), did not alter sGAG per DNA, and reduced total collagen per DNA content (Fig. 5c, Extended Data Fig. 10a). Notably, alcian blue staining revealed a large, rounded cell morphology and increased pericellular sGAG staining in response to loading, especially in early and reversed groups where loading was applied immediately after encapsulation, suggesting load-induced pericellular matrix (PCM) deposition. To test this we immunostained for PCM-exclusive collagen VI and found that all loaded groups exhibited increased collagen VI at the cell periphery, particularly in the groups loaded immediately, while free swelling controls were nearly devoid of ColVI (Fig. 5c). Message level expression of Col6a1 was significantly increased in early and delayed groups, suggesting that mechanical load is needed to initiate and maintain Col6a1 message level expression, resulting in ColVI accumulation in groups loaded immediately. Collagen VI is prevalent in the PCM of articular chondrocytes, but not in bone cell PCM, and functions to for resisting cellular deformation during cartilage matrix compression,^42^ mediating load-induced proliferation and chondrogenic gene expression.^43^ Thus, immediate loading, prior to significant matrix deposition, may have caused large cellular deformation, inducing production of a protective PCM and promoting chondrogenic differentiation. To that end, Sox9 message-level expression increased in all loaded groups, but was most significant when applied continuously (early; Fig. 5f), indicating a maintenance of chondrogenic gene expression that may explain the cartilage persistence seen *in vivo* in response to early loading. Similar trends were observed for Sox9-target genes ACAN and Col2a1 (Extended Data Fig. 10b). Consistent with the downregulation of YAP and downstream target CYR61 observed with TGF-β1-mediated chondrogenesis (Fig. 1), these genes are were also significantly decreased by chondrogenic loading (Extended Data Fig. 10b). Loading significantly decreased Col10a1 and OPN expression in groups where load was being applied at harvest (early and delayed), but not in the reversed group, suggesting load-mediated maintenance of the chondrogenic lineage (Fig. 5d). VEGF is an important factor for vascular invasion during endochondral healing, and is necessary for remodeling of the fracture callus. VEGF increased in all loaded groups but was most significant when applied after a period of free swelling (delayed), consistent with the effects of *in vivo* delayed loading enhanced endochondral bone growth (Fig. 5d), and suggests that loading may also regulate angiogenesis and vascular invasion.

**Figure 5:**
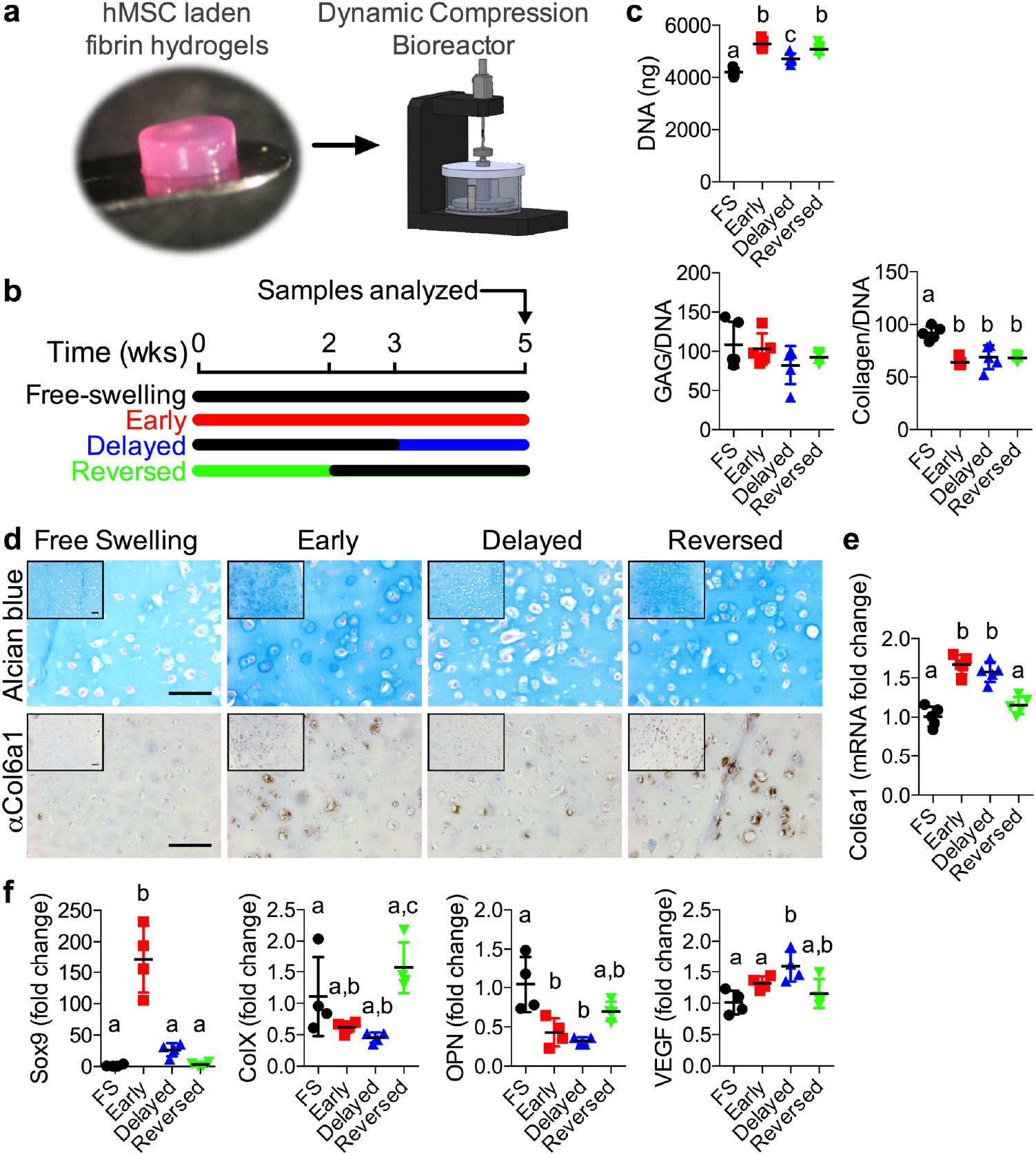
*In vitro* analysis of mechanical load on chondrogenic lineage progression. **a**, Dynamic compression was applied to hMSC-laden hydrogels (P4, 600,000 cells, 50mg/mL fibrinogen) in a custom-made bioreactor. Load was applied 2 hours per day, 5 days per week at 1 Hz and 10% strain after a 0.01 N preload was applied. **b**, Four loading groups were evaluated: free-swelling (FS) controls, early loading (continuous for 5 weeks), delayed loading (free-swelling for 3 weeks followed by 2 weeks of loading), and reversed loading (loading for 2 weeks followed by 3 weeks free-swelling). All samples were collected at week 5 for analysis. **c**, Quantification of biochemical DNA, sGAG, and total collagen content (n = 5). **d**, Alcian blue staining of load-induced sGAG concentrated around the cell periphery. **e**, Load induced pericellular Col6a1 deposition. Images shown at 20x with 10x insets, scale bars 100μm, n=2. **f-g**, Quantitative PCR at week 5 (n=4-5) showed significant load-dependent induction of (n = 4-5 per group) of Col6a1 (**f**) and Sox9, Col10a1, OPN, and VEGF (**g**). Relative expression was calculated as fold change over free swelling controls. Data are shown as mean ± s.d. with individual data points. Statistical comparisons between groups for each measure were performed by one-way ANOVA with Tukey’s *post-hoc* analyses, where groups sharing a letter (a,b,c) are not statistically different.

Angiogenesis enhances and accelerates the initiation of ossification in endochondral bone formation,^44–46^ and, as we showed previously, is influenced by mechanical cues during bone defect repair^6^ and can be modulated by scaffold architecture.^47^ To determine the influence of mechanical load timing on neovascularization during endochondral regeneration, we performed an additional study to quantify vascular invasion of the cartilage anlage in response to mechanical loading by microCT angiography. This was accomplished by perfusing the vasculature with a lead chromate-based contrast agent (Microfil MV-122) to attenuate X-rays in the vasculature^48–50^ for 3D quantification of blood vessel networks in and around the defect (Fig. 6, Extended Data Fig. 11). Sequential microCT scanning of the perfused limbs before and after bone decalcification enabled independent quantification of bone formation and vascularization in the same samples.^6^ In this study, we performed microCT angiography three weeks after the onset of loading for both early (n = 10) and delayed (n = 8) loading, with each animal assigned a loaded limb and a contralateral stiff plate control. Thus, animals in the early loading group (including their contralateral controls) were perfused at week 3 and the delayed group at week 7 (Fig. 6a,i). These time points were selected to capture the transient vascular network response to the dynamic mechanical environment.^6,46^ In the 5mm defect ROI, early loading did not alter bone formation at week 3 (Fig. 6b,d), consistent with the independent prior results at week 4 (cf. Fig. 2e), but significantly inhibited vascular ingrowth, blunting the preferentially axial orientation of the vessel network observed in the stiff group and producing a more isotropic distribution of orientations (Fig. 6c,e-h). In contrast, delayed loading enhanced bone formation at week 7 (Fig. 6j,l), consistent with the independent prior results at week 8 (cf. Fig. 2). However, delayed loading did not alter vascular morphometry parameters other than reduced vessel anisotropy (Fig. 6k,m-p). Loading did not alter the vascular structures of the peripheral muscle (7mm ROI), indicating a local effect of loading on either vessel recruitment by the anlage, endothelial cell invasion, or neovessel integrity (Extended Data Fig. 11). Together, these observations that mechanical loading did not affect the vascular supply in the peripheral musculature but decreased the degree of axial orientation of the invading vasculature suggest that loading may differentially regulate two distinct sources of angiogenic vessels, one from within the cortex, and one from the surrounding musculature. Loading may therefore disproportionally impair or delay the vascular invasion from the endocortical space, while allowing transverse invasion from the surrounding musculature.

**Figure 6:**
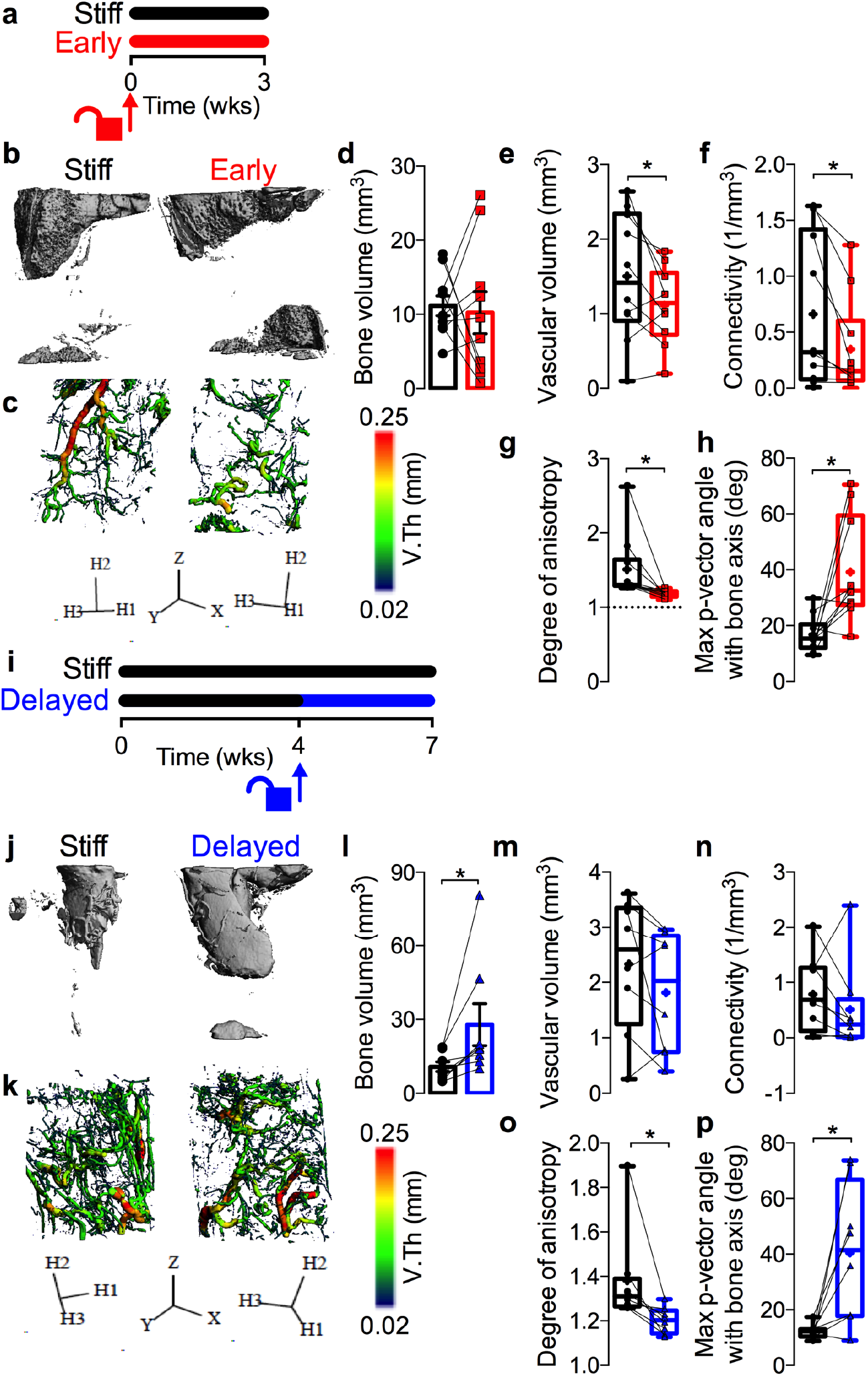
Mechanical control of neovascularization. Changes in vascular network morphometry were measured by microCT angiography in two independent cohorts of animals receiving either early or delayed loading, with each experimental limb paired with a contralateral stiff plate control (standard 5mm diameter defect ROI). **a**, Schematic of loading timeline with early loading featuring compliant plate unlocking at implantation and contrast agent perfused at week 3. **b-c**, Representative microCT reconstructions of bone (**b**) and blood vessels with local vessel diameter mapping (**c**) under stiff and early loading conditions at week 3, n = 10. **d**, Quantification of bone volume at week 3. **e-h**, 3D vascular network morphometry quantifying vascular volume (**e**), connectivity (**f**), and vessel orientation and distribution, as measured by degree of anisotropy (**g**) and the angle with respect to the bone-axis of the maximum principal eigenvector (H2) of the mean intercept length (MIL) tensor (**h**), indicating the dominant direction of vessel orientation. Degree of anisotropy represents the ratio of the longest and shortest MIL eigenvalues. DA=1 (dotted line) indicates isotropy. **i**, Schematic of loading timeline with delayed loading featuring compliant plate unlocking at week 4 and contrast agent perfused at week 7. **j-l**, Quantification of bone volume at week 7 with representative images. **m-p**, Vascular network morphometry measured by vascular volume (**m**), connectivity (**n**), degree of anisotropy (**o**), and maximum principal vector angle (**p**). All representative images were selected to best match the average microCT vascular morphometry of that group at the corresponding time point. Paired data shown either as mean ± s.e.m. or superimposed on box plots displaying median as horizontal line, inter-quartile range as boxes, and min/max range as whiskers. Mean values are indicated by +. Comparisons between groups evaluated by paired two-tailed Student’s t-tests (*p<0.05).

In development, the cartilage anlage initiates as an avascular template that, upon chondrocyte hypertrophy and calcification, is invaded by blood vessels from the surrounding tissue. To simultaneously quantify 3D cartilage and vascular tissue distribution in the newly formed defect tissue, we performed dual contrast agent-enhanced microCT imaging by cartilage contrast agent (CA^4+^) equilibration^51^ in samples already perfused with vascular contrast agent. CA^4+^ partitions at equilibrium with the negatively charged sulfated glycosaminoglycans (sGAG) and attenuates X-rays proportional to sGAG concentration.^51^ We then performed a region-of-interest (ROI) analysis, evaluating the vasculature and the cartilage in two regions: the defect core (1.5mm diameter) and in the surrounding annular region (5mm outer diameter). As hypothesized, the core region remained distinctly avascular in all groups (Figure 7a,b,d,e), though the sGAG-positive tissue distribution was not significantly different between regions (Figure 7c,f,g,h). This avascular core was not observed at the same time points in prior studies with BMP-2 delivery, in which bone formation occurred primarily through intramembranous ossification,^6^ suggesting this pattern of vascular exclusion may be specific to endochondral repair. Differences in cartilage volume between the stiff plate and early loading groups in annular and core regions did not reach statistical significance for either loading regime (Fig. 7c,f).

**Figure 7:**
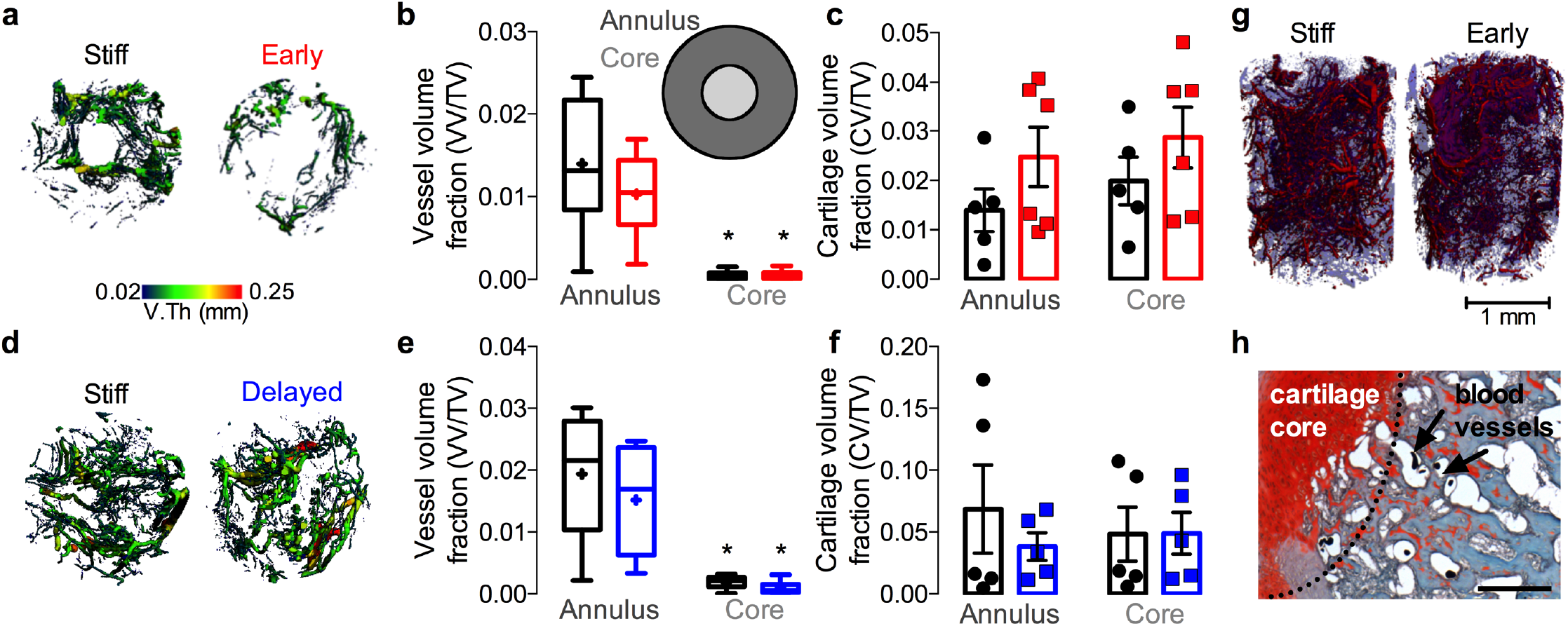
Spatial distribution of neovascular and cartilaginous tissues. Dual contrast agent-enhanced microCT imaging was performed three weeks after the onset of loading (at week 3 for early and week 7 for delayed) in comparison to contralateral stiff plate controls. **a**, Axial view of 3D neovessel diameter mapping under stiff and early loading conditions at week 3, n = 10. **b**, Region of interest analysis to quantify vascular volume fraction in a 1.5mm diameter core region compared to a 5mm-1.5mm annular region (inset). **c**, Cationic (CA^4^+) cartilage contrast agent-enhanced microCT quantification of cartilage in annulus and core regions. n = 5-6. **d**, Axial view of 3D neovessel diameter mapping under stiff and delayed loading conditions at week 7, n = 8. **e**, Region of interest analysis of vascular volume fraction. **f**, Cartilage contrast agent-enhanced microCT quantification of cartilage in annulus and core regions at week 7, n = 5-6. **g**, Representative image of co-registered contrast agent-enhanced cartilage imaging with microCT angiography of neovasculature where cartilage is shaded blue vessels are red. **h**, Safranin-O/Fast green-stained histological sections of vascular contrast agent-perfused tissues showing the avascular cartilage core and blood vessels in the surrounding tissue. Residual contrast agent exhibits thermal contraction during paraffin processing, visible as dark dots in vessel lumens. All representative images were selected to best match the average microCT-quantified cartilage morphometry of that group at the corresponding time point. Scale bar, 100 μm. Data shown either as mean ± s.e.m. with individual data points or with box plots displaying median as horizontal line, inter-quartile range as boxes, and min/max range as whiskers. Mean values on box plots are indicated by +. (*p<0.05, two-way ANOVA with Tukey’s *post-hoc* comparisons)

Vascularization and anlage maturation are linked at both cellular and molecular scales, and influence one another during endochondral ossification. For example, inhibition of angiogenesis can promote phenotypic stability of MSC-derived chondrocytes *in vivo*,^52^ while chondrocyte hypertrophy is in part responsible for neovessel recruitment.^1^ Thus, these data can be explained either by direct inhibition of axial blood vessel integrity, and cartilage maturation in turn, or inversely, by load-induced delay of chondrocyte hypertrophy and subsequent neovessel recruitment.

In this work, biopolymer, biodegradable microspheres capable of locally and temporally providing soluble chondroinductive cues^14,35,53^ were incorporated into hMSC condensations to provide the signals necessary to drive cartilage formation immediately upon implantation in femoral defects while not interfering with critical cell-cell interactions required for this process. The mechanical environment of these defects was then modulated via fixators with tunable compliance to promote the progression of endochondral ossification. Upon transplantation, *in vivo* mechanical loading regulated endochondral ossification and restored mechanical function to challenging bone defects (>60% larger than critical size). These findings demonstrate the importance of *in vivo* mechanical stimuli for functional, vascularized tissue regeneration through endochondral ossification of transplanted cells and implicate mechanical cues as important factors for successful biomimetic tissue engineering of other tissues whose form and function is dictated by mechanical stimuli during development and homeostasis.

## Limitations

These studies used athymic (RNU) rats to facilitate xenogenic cell transplantation and assessment of functional hMSC engraftment,^25^ but this model may miss some immunomodulatory functions of the transplanted cells.^54^ However, potential translational application of these findings to the clinic would likely involve autologous cell transplantation, which would not illicit a T-cell response, or immune-matched allogeneic cells. This makes the RNU a reasonable model for this approach.^25^ Additionally, the rnu/rnu strain maintains an intact thymus-independent immune response including natural killer cells,^54^ which are known to be inhibited by MSCs,^54^ suggesting that some of the immunomodulatory function may be maintained in this model. A detailed exploration of hMSC-immune cell interactions during endochondral bone regeneration has not been performed and warrants investigation, but is outside the scope of the present study.

The model used in these studies features bilateral defects, with compliant (early or delayed) and stiff plates distributed in either of two ways: 1) evenly distributed among limbs such that each fixation plate is paired with each other type in equal numbers, or 2) each compliant plate (early or delayed) is paired with a contralateral stiff fixation plate. This design is used for two reasons. First, it enables accounting for animal-animal variability and microfil perfusion quality in statistical models. Second, should preferential weight bearing cause reduced loading of the limbs that are affixed with compliant plates, this preferential unloading would be further exacerbated if the contralateral limb were unoperated. In prior experiments using this model, we have not observed systemic or contralateral effects of hMSC transplantation^25^ or growth factor presentation^32^ on either bone formation or angiogenesis^6,24^. In this study, the ipsilateral response did not depend on the experimental group of the contralateral limb. This suggests that ambulatory imbalance is low or not sufficient to alter load-induced bone regeneration. Daily observation of animal ambulation and *in vivo* x-ray videography^15,30^ during treadmill ambulation demonstrate that compliant plates do not impair weight bearing in either limb during ambulation. Further, we previously evaluated gait kinematics in response to fixation plate insertion and bone defect creation and found no significant differences between groups or compared to sham-operated controls, indicating that defect creation and fixation implantation does not significantly alter gait kinematics.^30^

One limitation of this study was our use of only one hMSC donor. A single donor was chosen to reduce the compounding factor of donor variability. However, this MSC/TGFb aggregate sheet system has been used previously with additional donors to heal calvarial defects with great success,^35^ indicating the robustness of the system regardless of donor.

While we have shown that this developmentally inspired approach reproduces aspects of endochondral ossification and results in functional bone in critically-sized defects, it does not fully recapitulate the intricate and tightly coordinated process of limb development. Currently our approach is limited to one potent developmental morphogen and a mechanical strain environment dependent on tissue composition. Future studies will be needed to more accurately represent the biochemical and mechanical environment seen throughout mesenchymal condensation and limb bud endochondral ossification.

### Methods

Methods are available in the online version of the paper.

### Competing interests

The authors declare they have no competing financial interests.

## Acknowledgements

We thank the staff of the Freimann Life Science Center (FLSC) at the University of Notre Dame for animal care and husbandry, the staff of the Notre Dame Integrated Imaging Facility (NDIIF) for imaging support, and Amad Awadallah of the Case Western Reserve University Histology Core Facility for technical support. We thank Glen Niebur, University of Notre Dame, for insightful comments on the manuscript. We gratefully acknowledge funding from the Naughton Foundation (A.M.M., J.D.B.), the Indiana Clinical and Translational Sciences Institute, grant number UL1TR001108 from the National Institutes of Health (J.D.B), the American Heart Association, grant number 16SDG31230034 (J.D.B), the National Science Foundation, grant number 1435467 (J.D.B), the National Institutes of Health’s National Institute of Arthritis and Musculoskeletal and Skin Diseases under award numbers R01AR066193 (E.A), R01AR063194 (E.A.), and R01AR069564 (E.A.), the National Institutes of Health’s National Institute of Biomedical Imaging & Bioengineering under award number R01EB023907 (E.A.), the National Institutes of Health’s National Institute of Dental and Craniofacial Research under award number 5F32DE024712 (S.H.), and the Ohio Biomedical Research Commercialization Program under award number TECG20150782 (E.A.). The contents of this publication are solely the responsibility of the authors and do not necessarily represent the official views of the National Institutes of Health, the National Science Foundation, or other funding agencies.

## Author contributions

E.A. and J.D.B. conceived and supervised the research. A.M.M., S.H., E.A., and J.D.B. designed the experiments, analyzed the data, and wrote the paper. All authors collected data, and commented on and approved the final manuscript. Correspondence and requests for materials should be addressed to E.A. or J.D.B.

## Extended Data

**Extended Data Figure 1:**
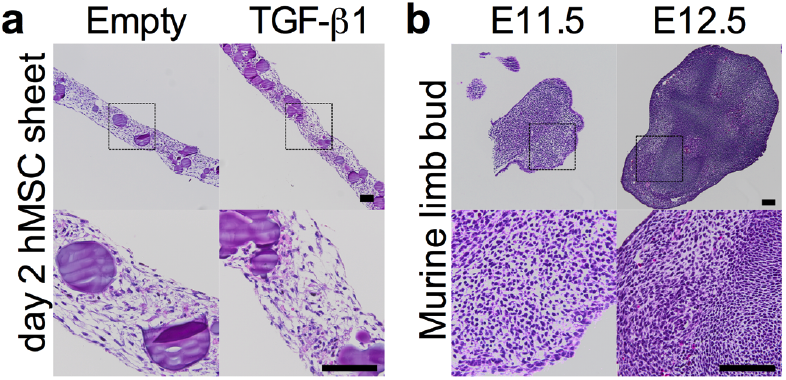
H&E staining of mesenchymal condensations and murine embryonic limb buds. **a**, Haematoxylin and eosin (H&E) staining of hMSC sheets cultured for two days on transwell inserts with empty gelatin microspheres (left), or loaded with 600 ng TGF-β1 (right). Bottom: magnification of enclosed areas. **b**, H&E staining of murine limb buds at embryonic days 11.5 and 12.5 (E11.5, E12.5). Bottom: photomicrographs of embryos with limb buds indicated by arrows. Scale bars, 100 μm. Histological images are representative of 3 independent samples per group.

**Extended Data Figure 2:**
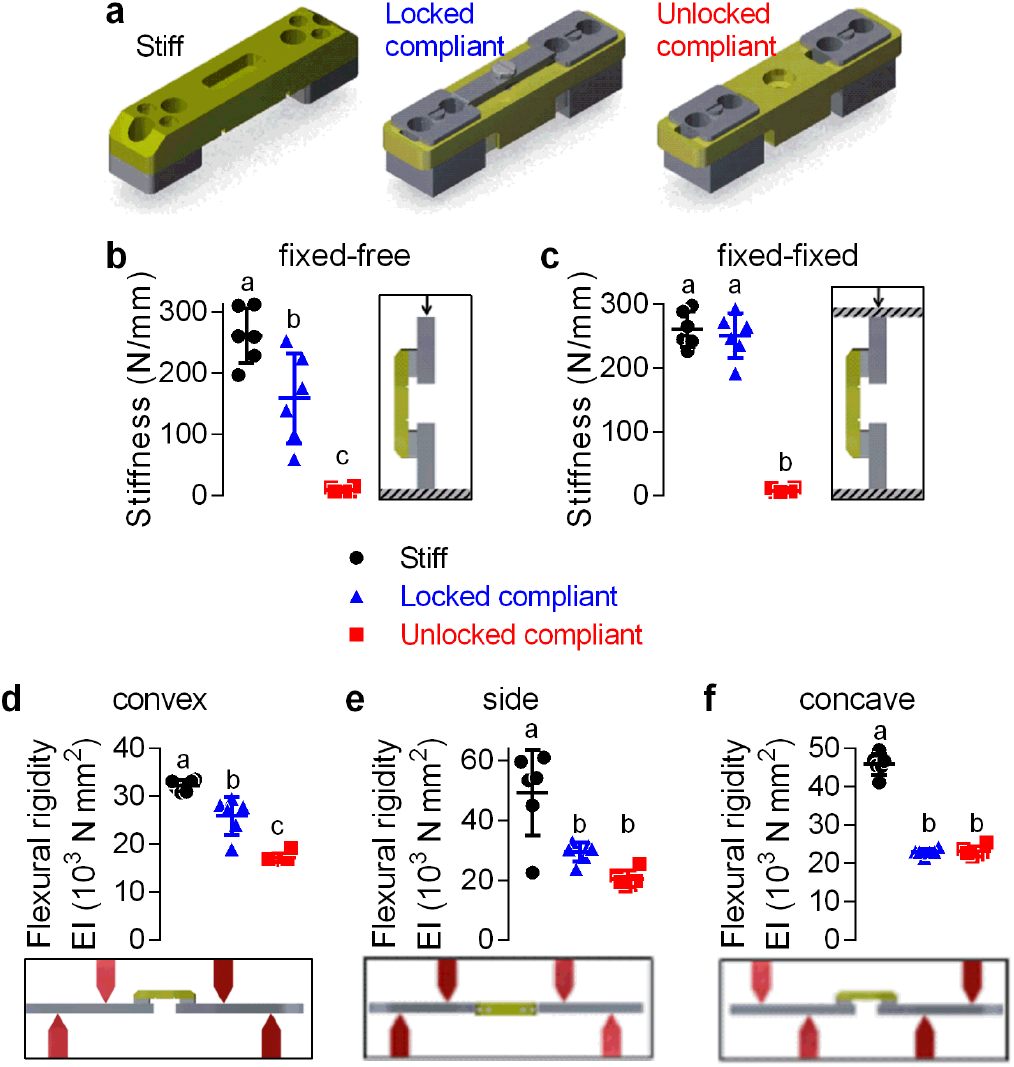
Fixation plate characterization. **a**, Bilateral critically-sized (8 mm) defects in the femora of 14 week-old, male RNU rates were stabilized by fixation plates that enabled dynamic control of ambulatory load sharing (n = 6 per group). **b-c**, Axial compression testing with fixed-fixed or fixed-free boundary conditions, as illustrated. **d-f**, Four-point bend testing in three directions, as illustrated, with supports and load applied to maintain constant bending moment on the plate. Statistical comparisons evaluated by oneway ANOVA on log-transformed data to ensure normality of residuals followed by Tukey’s *post hoc* test. Summary data shown as individual data points with mean ± s.d.

**Extended Data Figure 3:**
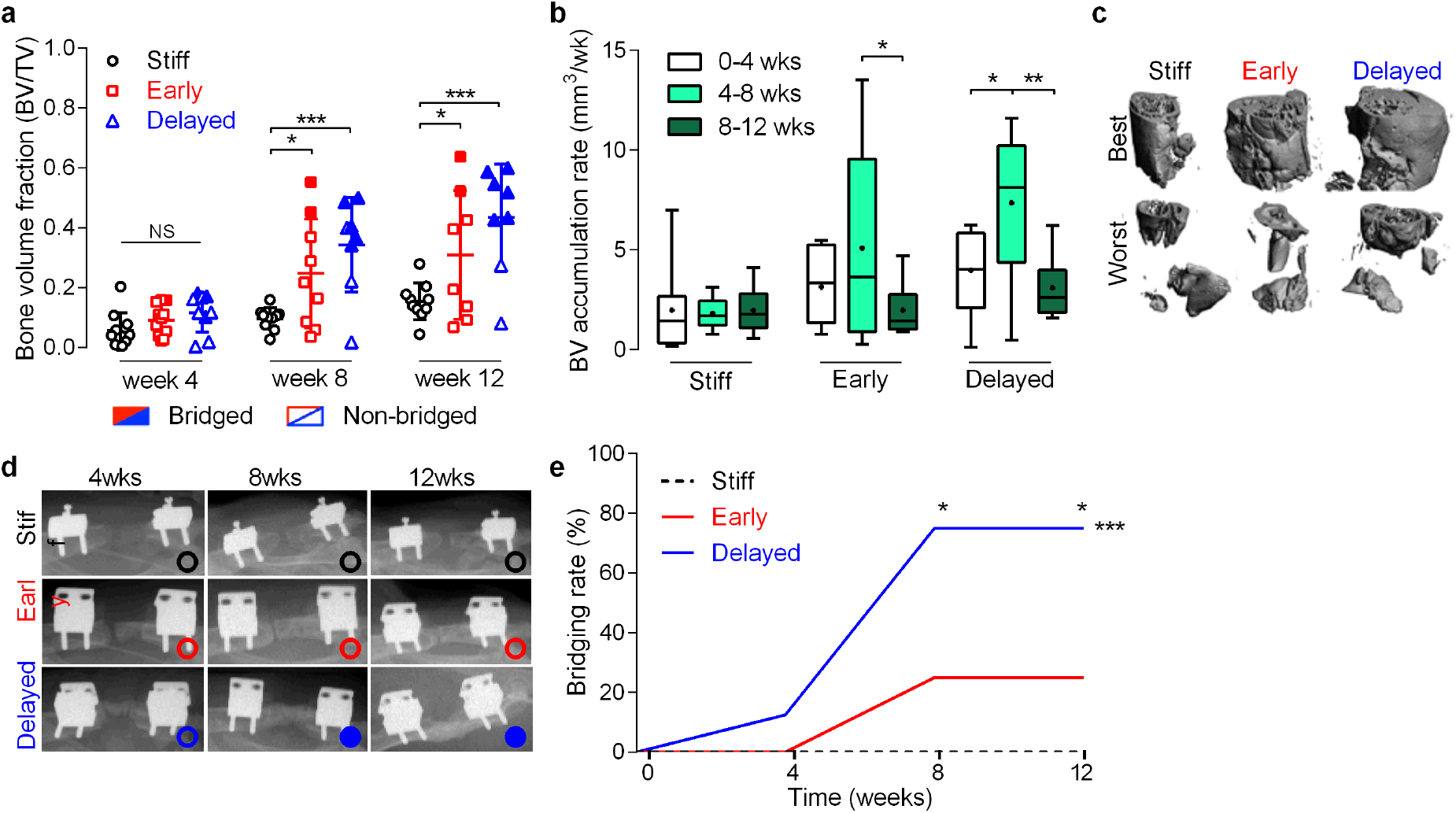
Bone accumulation and bridging rates. **a**, Bone volume fraction (BV/TV) over time where bone volume is normalized to a standard total volume of the defect area (8mm length × 5mm diameter). Individual data points shown with mean ± s.d. Samples with bony bridging are shown in shaded data points, while open data points indicate non-bridged. **b**, accumulation rate, defined as bone volume accrual over each 4-week interval. Box plots display median as horizontal line, inter-quartile range as boxes, and min/max range as whiskers. Mean values are indicated by +. BV accumulation rate was elevated in both early and delayed groups between weeks 4 and 8. *p<0.05, **p<0.01, ***p<0.001, two-way ANOVA with Tukey’s *post-hoc* analysis. **c**, High resolution (20 μm voxel size) MicroCT reconstructions of excised femurs at week 12 showing best- and worst-case regeneration for each group. **d**, Representative x-ray images for each group at 4, 8 and 12 weeks illustrating bridged (filled circles) and non-bridged (open circles) samples; images chosen based on mean bone volume at week 12. **e**, Longitudinal analysis of bone bridging *in vivo* at 4 (n= 11, 11, 9, for stiff, early, and delayed, respectively), 8 (n =10, 9, 8) and 12 weeks (n = 10, 8, 8) determined via x-ray as mineral fully traversing the defect. Significance of trend was analyzed by chi-square test for trend (***p<0.001) while differences between groups were determined by chi-square test at each time point with Bonferroni correction (*p<0.05). All stiff group samples failed to bridge by week 12, while 25% and 75% of the early and delayed loading groups achieved bridging by this time point, respectively.

**Extended Data Figure 4:**
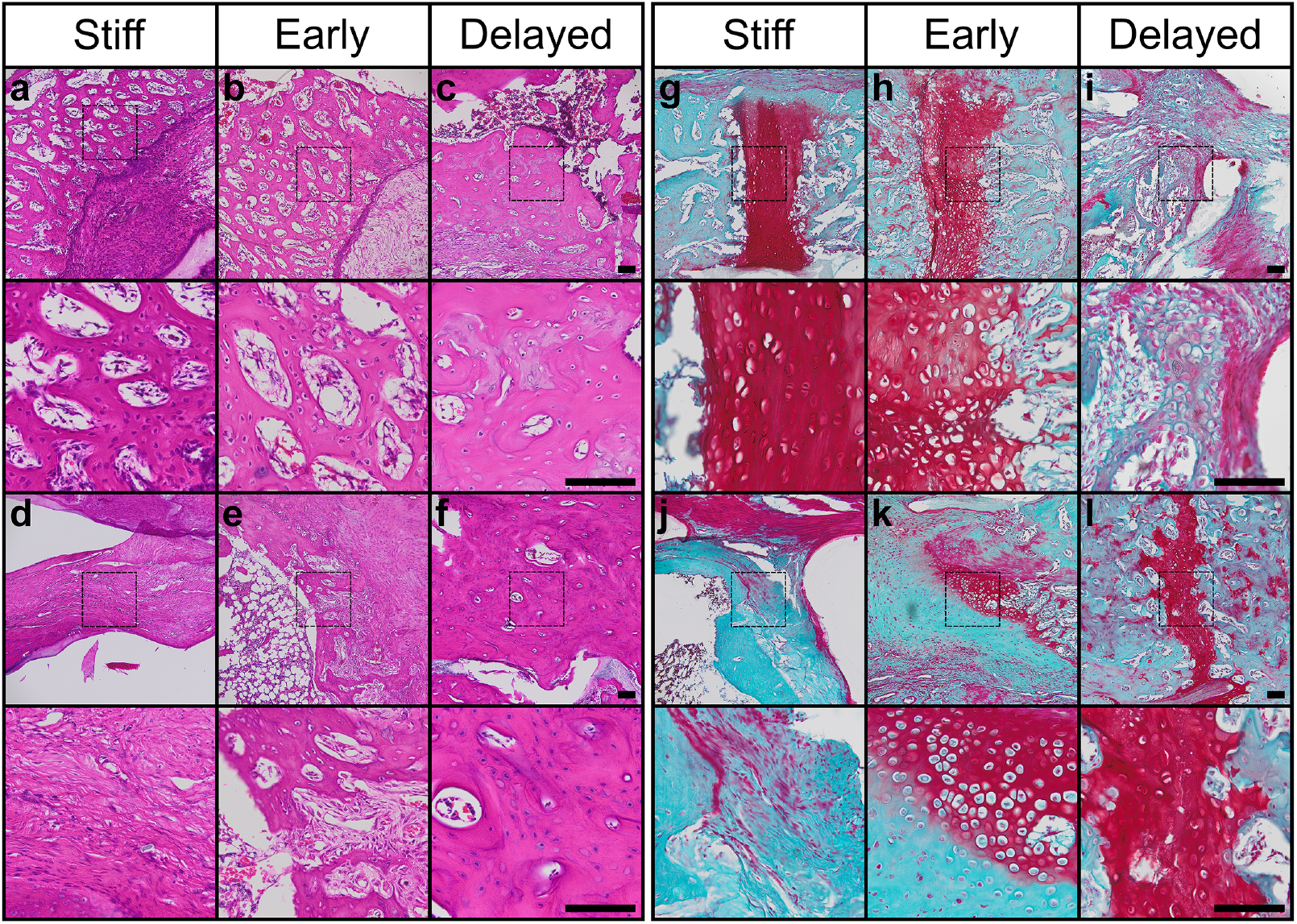
Full histological analysis of endochondral bone formation. **a-f**, H&E staining. g-I, Safranin-O/Fast green staining at 10x (top) and 40x (bottom, magnification of dotted squares). Scale bars, 100 μm. Panels **a-c** and **g-I** show tissues at week 4 and panels **d-f** and **j-l** show tissues at week 12 (n = 1 per group at each time point, chosen by proximity to mean bone volume *in vivo* at 4 and 12 weeks). Bone formation in the defect exhibited porous trabecular morphology, though stiff and early groups exhibited some fibrotic and fibrocartilaginous tissue at week 12. Safranin-O staining revealed that non-mineralized bands observed in microCT reconstructions consisted of mature and hypertrophic chondrocytes, giving rise to woven and neotrabecular bone. The stiff and early groups exhibited fibrocartilage and hypertrophic cartilage at week 12. The delayed loading group had remnants of hypertrophic growth plate-like cartilage that was fused and embedded in mineralized bone.

**Extended Data Figure 5.**
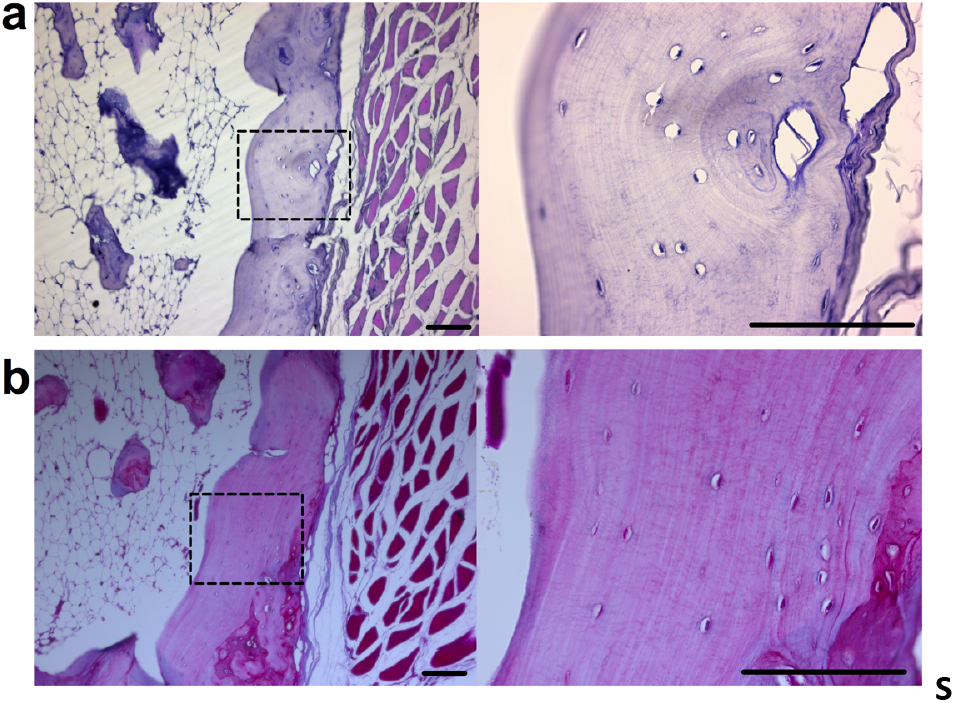
Representative histology of bone formation induced by 5ug of rhBMP2 delivered on an absorbable collagen sponge. **a**,**b**, Transverse tissue sections stained by H&E (**a**) and Safranin-O/Fast green (**b**) at 10X and 40X . 40X images are magnifications of the dotted squares in the respective 10X images. Scale bars, 100um.

**Extended Data Figure 6:**
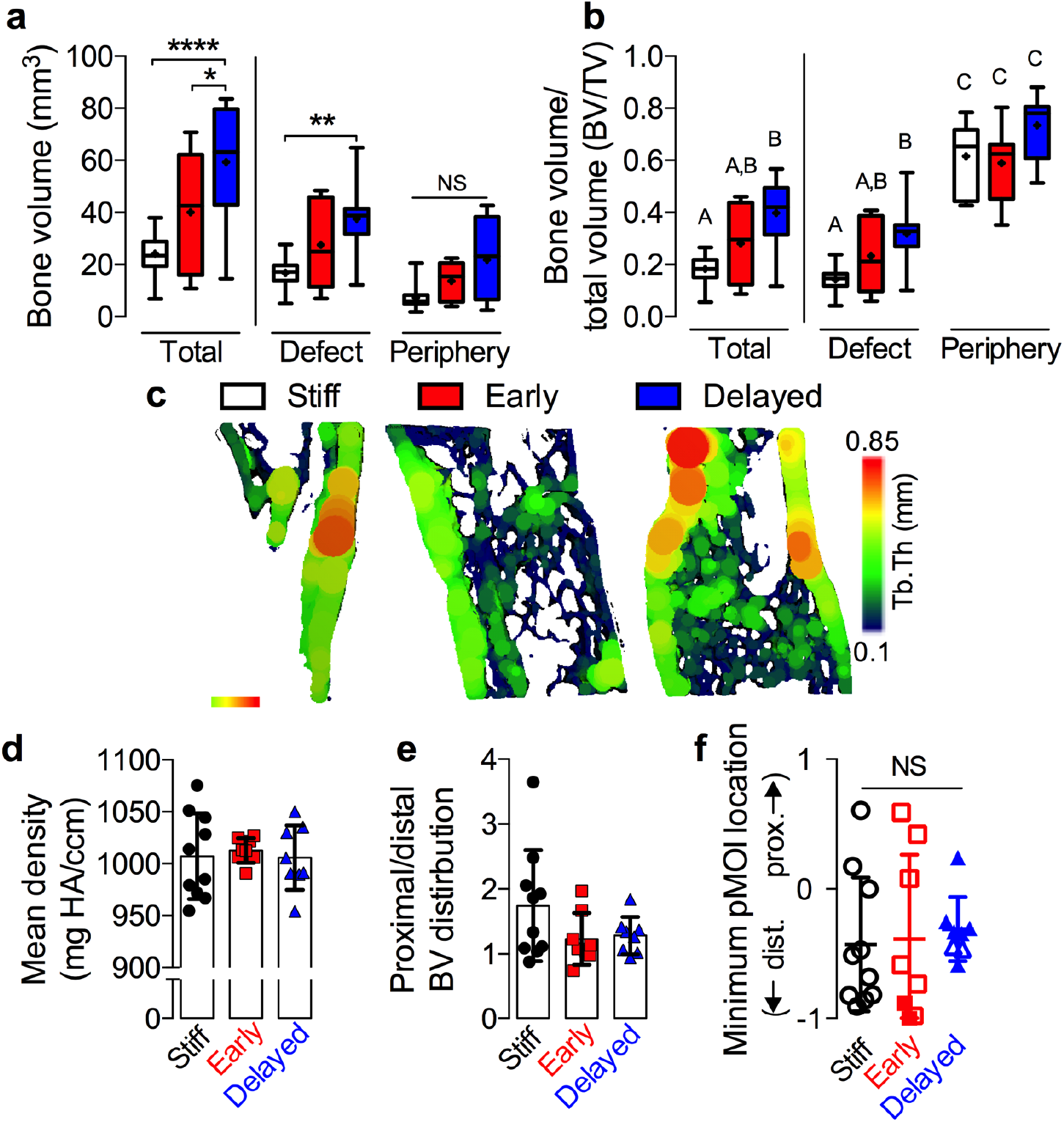
MicroCT region of-interest analysis demonstrating formation of cortical and trabecular bone compartments. **a**, Region of interest (ROI) analyses of bone volume in total, defect, and periphery ROIs, defined by regions either inside a 5mm diameter cylinder (defect) or annulus with 7 mm outer diameter and 5 mm inner diameter (periphery, n = 10, 8, 8 for stiff, early, and delayed respectively). Delayed mechanical loading enhanced bone formation, especially in the defect ROI. **b**, ROI analysis of bone volume fraction. Groups with shared significance indicator letters have no significant differences. Bone was most concentrated in the ectopic region, indicative of a cortical shell, but differences between groups were most prominent in the defect region. Increased thickness in a cortex-like shell was evident in each group. **c**, Representative microCT images of sagittal sections with mean trabecular thickness mapping overlay; images selected based on mean bone volume. **d**, Mean mineral density. No differences between groups were detected. **e**, Proximal vs. distal region of interest analysis, expressed as ratio of bone volume in proximal to distal halves of the defect. Data showed a trend toward greater bone formation in the proximal half of the defect, especially in the stiff plate group. **f**, The location of minimum pMOI for each sample, with mean ± s.d. In panel **f**, filled data points indicate bridged samples and open data points indicate non-bridged samples. Box plots show interquartile range with whiskers at minimum and maximum values, center lines at median, and + symbols at the mean. Bar graphs show data with mean ± s.d. *p<0.05, ** p<0.01, **** p<0.0001, NS = not significant, one or two-way ANOVA with Tukey’s *post-hoc* analysis.

**Extended Data Figure 7:**
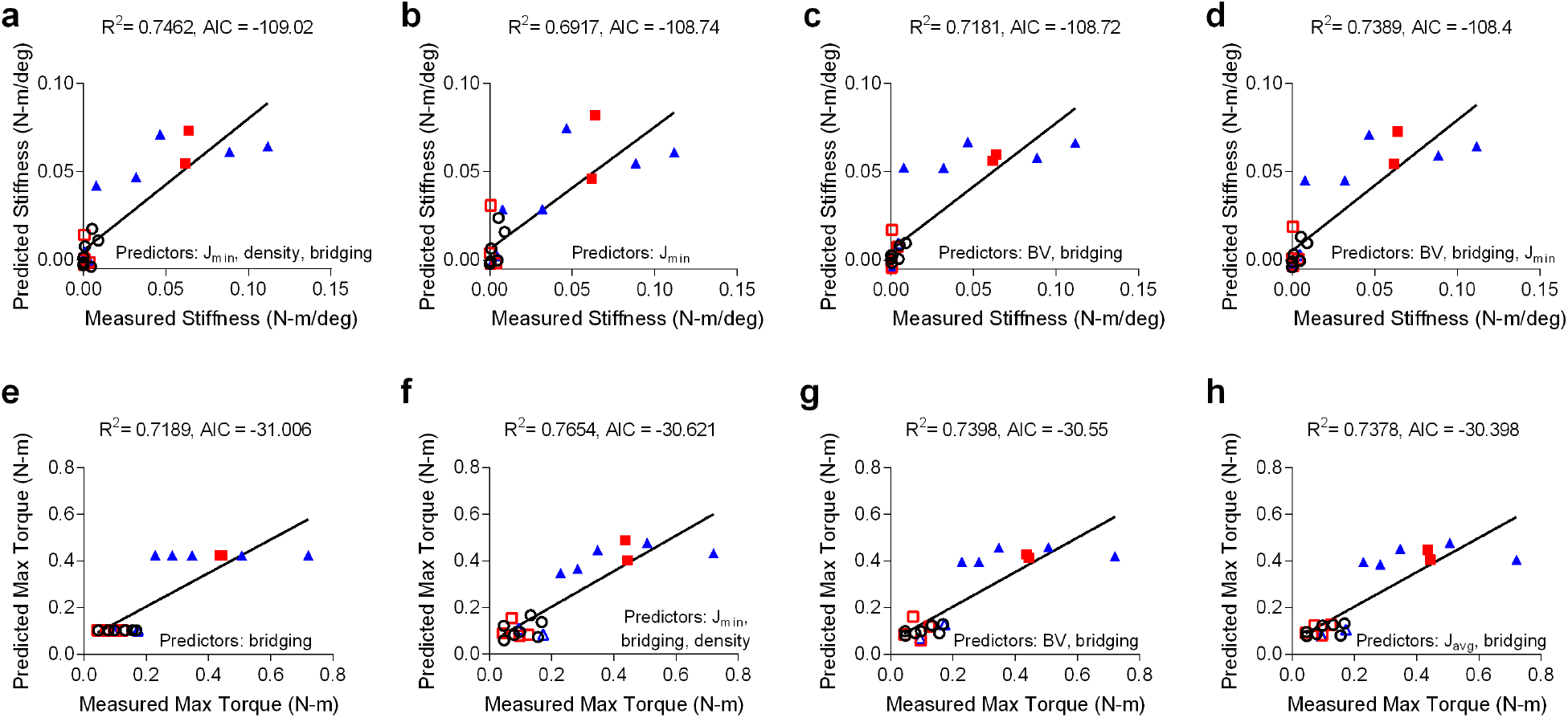
Best subsets analysis of mechanical testing data, models 2-5. The top 5 models, where best model selection was made by minimization of Akaike’s Information Criterion (AIC). Predictive models of **a-d**, torsional stiffness or **e-h**, maximum torque at failure were composed of combinations of the parameters average polar moment of inertia (J_avg_), minimum polar moment of inertia (J_min_), bone volume (BV), binary bridging score, and average mineral density. Type II regression was used to determine correlations between predicted and measured values. Shaded data points indicate bridged samples

**Extended Data Figure 8:**
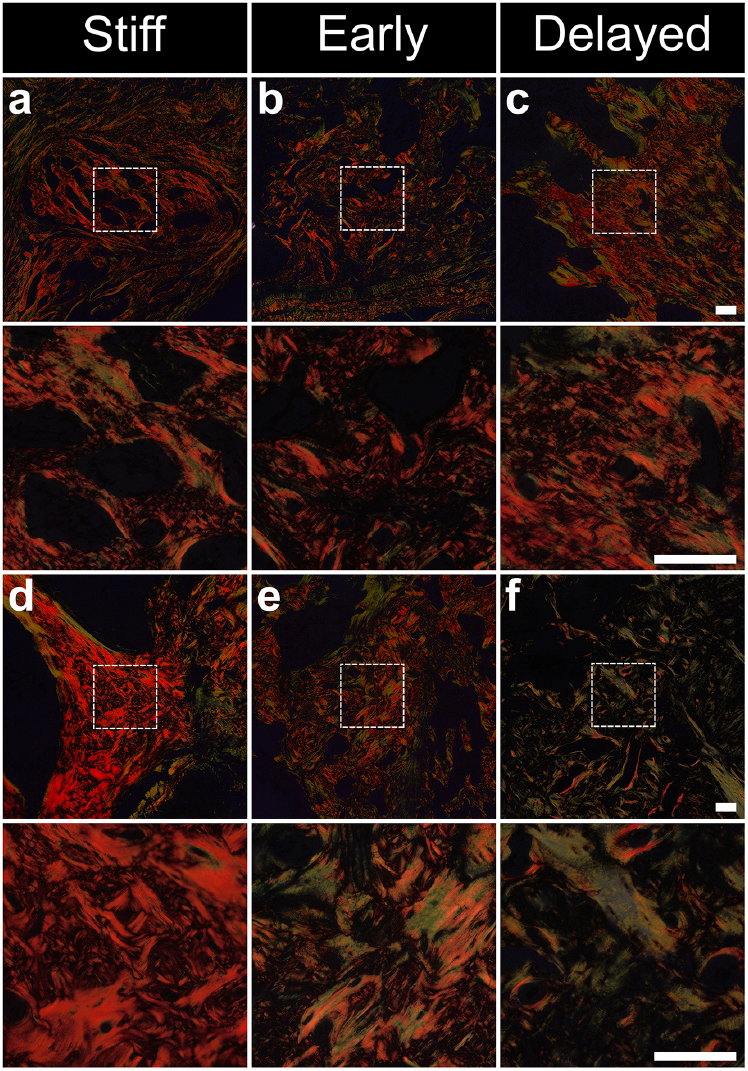
Collagen organization. Photomicrographs of picrosirius red staining obtained using polarized light microscopy on sections at week 4 (**a-c**) and 12 (**d-f**). No differences between groups were apparent at week 4, and were characterized primarily by woven bone, but by week 12, both early and delayed loading groups exhibited reduced collagen fiber birefringence compared to stiff controls, indicative of prolonged new bone formation or delayed remodelling to lamellar bone (n = 1 per group at each time point, chosen by proximity to mean bone volume *in vivo* at 4 and 12 weeks). Under polarized light, large collagen fibers birefringe yellow and orange, while thinner fibers are green. Images are shown at 10x (top) and 40x (bottom) magnification of dotted boxes. Scale bars, 100 μm.

**Extended Data Figure 9:**
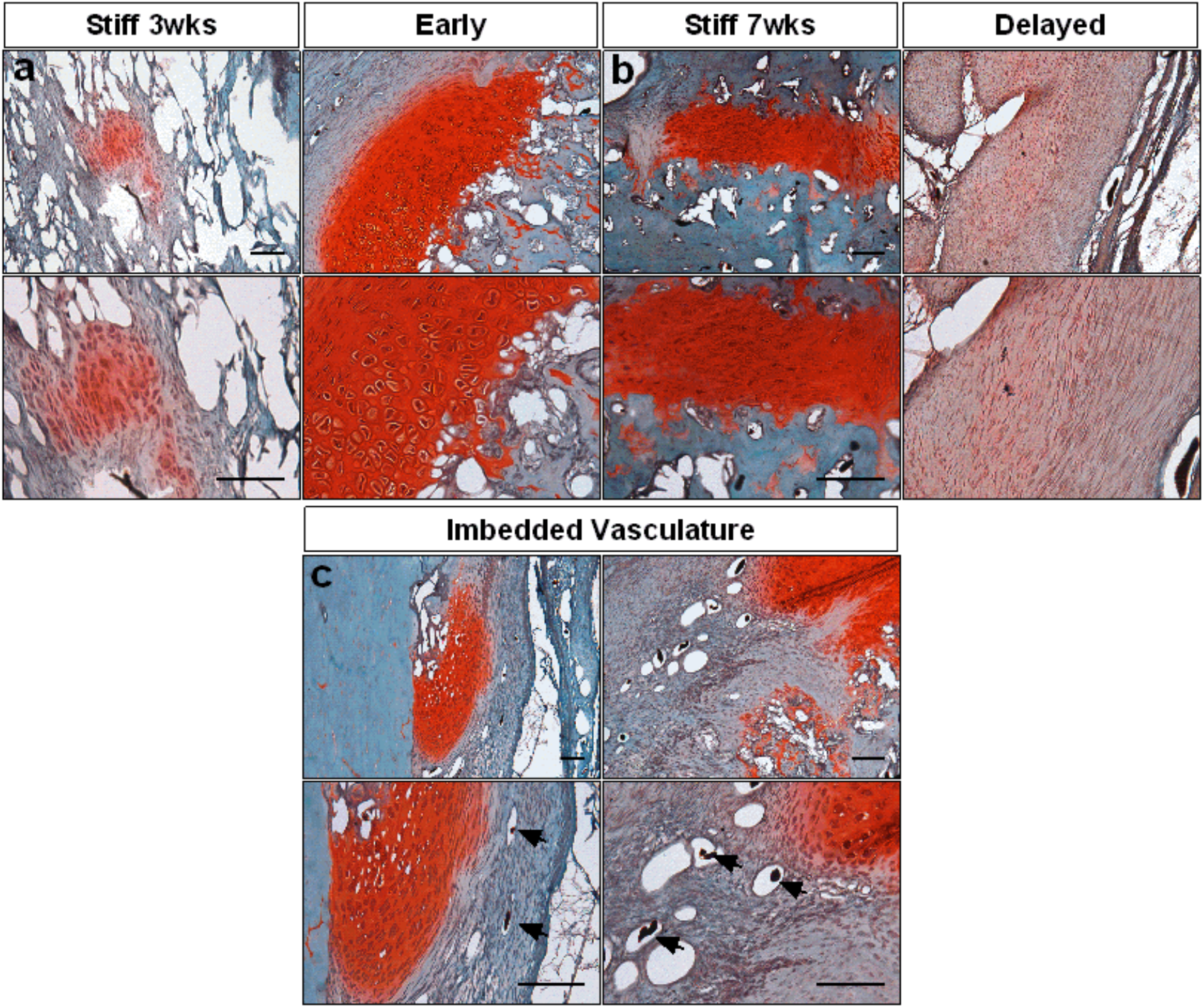
Histology of contrast agent-perfused sections. Safranin-O/Fast green staining at 10x (top) and 20x (bottom) of sagittal histological sections of defects at **a**, 3 weeks with early loading and **b**, 7 weeks with delayed loading, shown alongside stiff contralateral controls at either time point (n = 3 per group at each time point, chosen by proximity to mean vessel volume at 3 and 7 weeks) displaying increased cartilage with early loading. **c**, Additional Safranin-O/Fast green sections at 10x (top) and 20x (bottom) of early (left) and stiff (right) illustrate invading vascular imbedded in bone matrix adjacent to cartilage, indicative of endochondral ossification. Patent vessels can be identified by residual perfused contrast agent (arrows). Scale bars, 100 μm.

**Extended figure 10:**
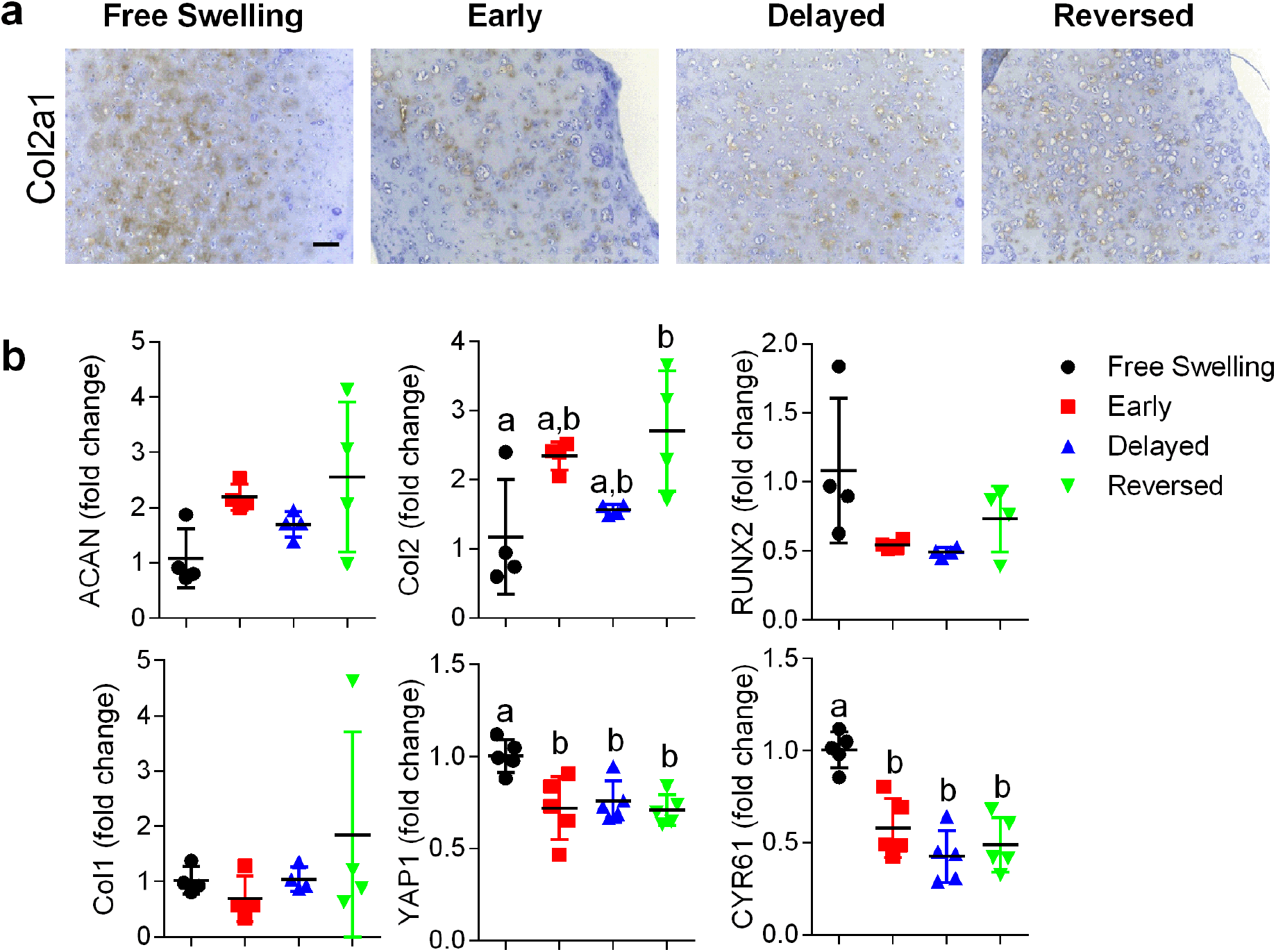
Message level expression of dynamically compressed hydrogels. **a**, Col2a1 deposition in samples after 5 weeks *in vitro* with hematoxylin counterstain (10x images, scale bars 100μm, n=2). **b**, Additionally, samples were analyzed at 5 weeks for message level expression of ACAN, Col2a1, RUNX2, Col1a1, YAP1, and CYR61 via qPCR (n=4-5) calculated as fold change over free swelling controls. Data are shown as mean ± s.d. with individual data points. One-way ANOVA and multi comparison by tukey’s post hoc was used to determine significance (p< 0.05) where groups sharing a letter are not statistically different.

**Extended Data Figure 11:**
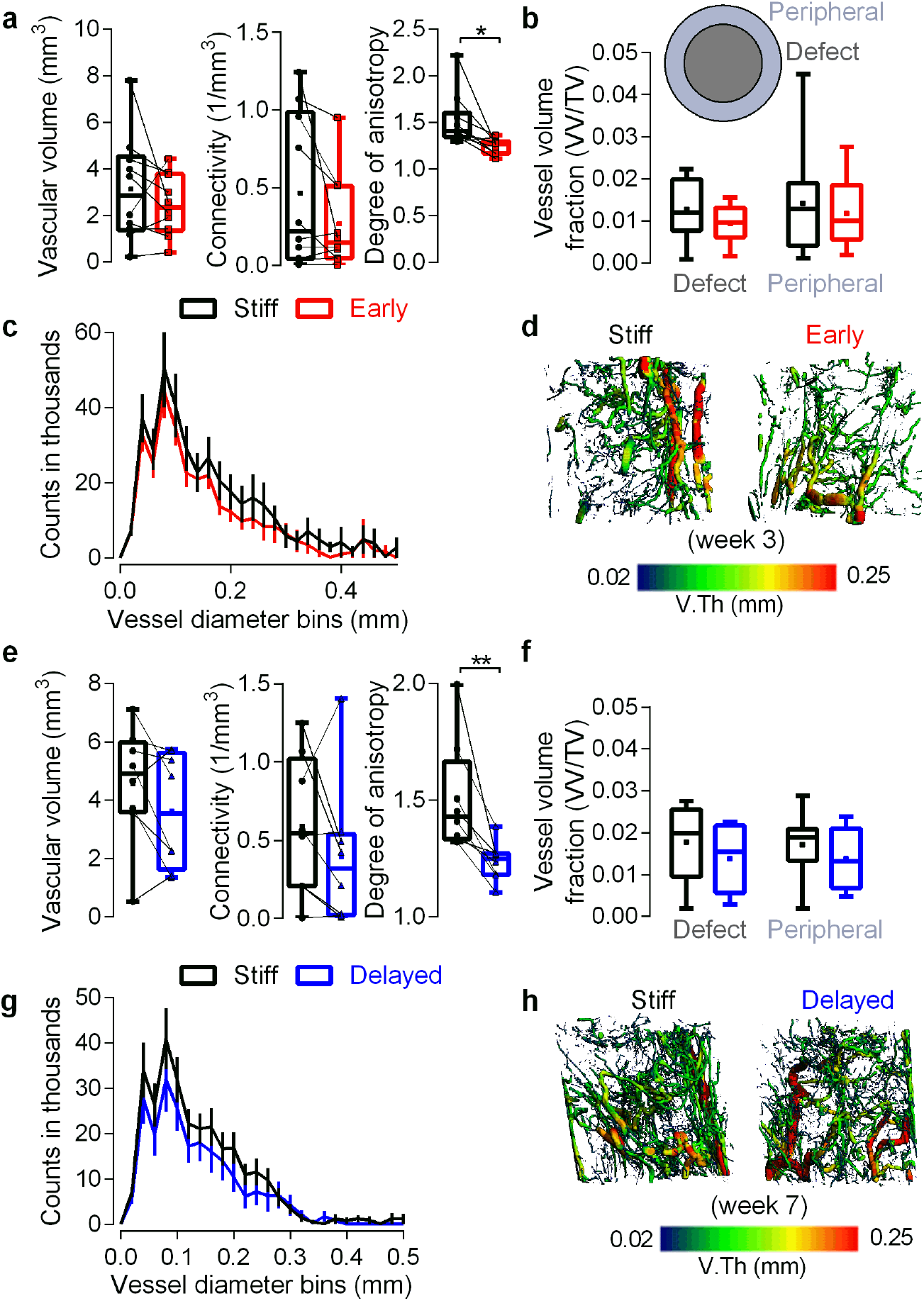
Vasculature in defect periphery. Vessel volume, connectivity, and anisotropy of **a**, early loaded limbs (n = 10) compared to contralateral stiff controls in a 7mm region of interest that included vasculature from peripheral muscle, where no differences were seen in any morphometric parameter in either group apart from vessel anisotropy (*p<.05, **p<0.01, two-way paired student’s t-test). Paired individual data points are superimposed on box plots displaying median as horizontal line, inter-quartile range as boxes, and min/max range as whiskers. Mean values are indicated by +. **b**, Region of interest analysis demonstrating no differences in vascular volume fraction in the 5mm diameter defect region compared to the 7mm-5mm peripheral region that included surrounding muscle (inset; two-way ANOVA with Tukey’s *post-hoc* comparisons). Box plots display median as horizontal line, inter-quartile range as boxes, and min/max range as whiskers. Mean values are indicated by +. **c**, Histograms of vessel diameter bins indicating similar vessel thickness distribution between groups, and **d**, representative microCT angiography vessel thickness mapping. Vessel volume, connectivity, and anisotropy of **e**, delayed loaded limbs (n = 8) compared to contralateral stiff controls in the same 7mm region where again no differences were seen in any morphometric parameter in either group apart from vessel anisotropy (*p<.05, **p<0.01, two-way paired student’s t-test). Data displayed as described previously. **f**, Region of interest analysis demonstrating no differences in vascular volume fraction in the defect region compared to the peripheral region (two-way ANOVA with Tukey’s *post-hoc* comparisons). Data displayed as previously described. **g**, Histograms of vessel diameter bins and **h**, representative microCT angiography vessel thickness mapping. All representative images chosen by proximity to mean vessel volume.

**Extended Data Video 1: Fixation plate actuation and bone regeneration.** Video animating plate configurations (top row) in which stiff plates minimize loading to the defect and compliant plates allow ambulatory load either immediately upon implantation (early) or after 4 weeks *in vivo* via surgical unlocking (delayed). Representative *in vivo* microCT reconstructions (bottom row) of regenerating defects at week 12, chosen by proximity to mean bone volume, demonstrate enhanced healing in the delayed group with complete bridging, while early and stiff groups failed to bridge by week 12, and stiff groups with devitalized constructs only achieved capping of bone ends.

## Materials and Methods

### hMSC isolation and expansion

Human mesenchymal stem cells (hMSCs) were derived from the posterior iliac crest of a healthy male donor (41 years of age) using a protocol approved by the University Hospitals of Cleveland Institutional Review Board. Cells were isolated using a Percoll gradient (Sigma-Aldrich, St. Louis, MO) and cultured in low-glucose Dulbecco’s modified Eagle’s medium (DMEM-LG; Sigma-Aldrich, St. Louis, MO) containing 10% prescreened fetal bovine serum (FBS; Sigma-Aldrich), 1% penicillin/streptomycin (P/S; Fisher Scientific), and 10 ng/ml fibroblast growth factor-2 (FGF-2, R&D Systems, Minneapolis, MN) as described previously.^14,53,55^ Cells were verified to be negative for mycoplasma contamination during expansion, prior to *in vivo* implantation.

### Gelatin microsphere synthesis and TGF-β1 loading

Gelatin microspheres were engineered and characterized as previously described.^55–57^ Briefly, an aqueous solution of 11.1% (w/v) acidic gelatin type A (Sigma-Aldrich) was added drop-wise to 250 ml of preheated (45°C) olive oil (GiaRussa, Coitsville, OH) and stirred at 500 rpm for 10 min. The solution temperature was then lowered to 4°C with constant stirring. One hundred milliliters of chilled acetone were added to the solution after 30 min and again after 1 h. The solution was then stirred at 1000 rpm for 5 min. The resulting microspheres were collected by filtration, washed with acetone to remove residual olive oil, and dried overnight at room temperature. Microspheres were then crosslinked for 4 h with 1% (w/v) genipin (Wako USA, Richmond, VA), washed 3 times with diH_2_O, and lyophilized. Hydrated GM gelatin microspheres were light blue in color and spherical in shape with an average diameter of 52.9±40.2 μm (N=150) and a crosslinking density of 25.5 ± 7.0% as determined by light micrograph image analysis (ImageJ software; National Institutes of Health, Bethesda, MD) and ninhydrine assay, respectively, according to protocols described previous.^33–34^ Growth factor-loaded microspheres were prepared by soaking crosslinked, UV-sterilized gelatin microspheres in a 80 μg/ml solution of rhTGF-β1 (Peprotech, Rocky Hill, NJ) in phosphate buffered saline (PBS) for 2 h at 37°C. To ensure 100% growth factor binding, the volume of TGF-β1 solution used was less than the equilibrium swelling volume of gelatin microspheres. Unloaded microspheres without growth factor were hydrated similarly using only PBS. TGF-β1 release from gelatin microspheres with comparable specifications in collagenase-containing PBS has been demonstrated to occur with an initial burst of ~60% followed by complete release of all loaded growth factor by 10 d.^57^

### Microsphere-incorporated hMSC sheet preparation

Expanded hMSCs (2×10^6^ cells/sheet; passage 4) were thoroughly mixed with TGF-β1-loaded gelatin microspheres (400 ng/mg; 1.5 mg/sheet) in serum-free, chemically defined medium comprised of DMEM-HG (Sigma-Aldrich) with 10% ITS^+^ Premix (Corning), 1 mM sodium pyruvate (HyClone), 100 μM non-essential amino acids (Lonza), 100 nM dexamethasone (MP Biomedicals, Solon, OH), 0.05 mM L-ascorbic acid-2-phosphate (Wako), and 1% P/S (Fisher Scientific) as described previously.^10^ Five hundred microliter of the suspension were seeded onto the pre-wetted membrane of transwell inserts (3-μm pore size, 12-mm diameter; Corning) using large orifice tips and allowed to self-assemble for 2 days. After 24 h, the medium in the lower compartment was replaced with 1.5 ml of fresh chemically defined medium. Control sheets incorporated with unloaded gelatin microspheres were prepared and cultured in a similar fashion. After 48 h, hMSC sheets embedded with TGF-β1-loaded gelatin microspheres were used for implantation. Sheets designated for *in vitro* analysis were harvested and cut in half at the midline using a razor blade.

### Surgical procedure

Critical-sized (8-mm) bilateral segmental defects were created in the femora of twenty 14 week-old male Rowett nude (RNU) rats (Charles River Labs, Wilmington, MA) under isoflurane anesthesia, as previously described.^26^ Briefly, anterolateral incisions were made over the length of each limb, and the vastus intermedius and vastus lateralis muscles were blunt-disected to expose the femur. Limbs were stabilized by custom internal fixation plates that allow controlled transfer of ambulatory loads *in vivo*^15^ and secured to the femur by four bi-cortical miniature screws (J.I. Morris Co, Southbridge, MA). The experimental design featured three groups. Control limbs (stiff) were stabilized with fixation plates that limited load transfer (n=16). Early loading limbs were stabilized by axially compliant fixation plates that allowed load transfer immediately upon implantation (early, n=12). Delayed loading limbs were stabilized by the same compliant plates initially implanted in a locked configuration to prevent loading, but after four weeks the plates were surgically unlocked to enable load transfer (n=12). Fixation plate mechanical characterization is detailed in Extended Data Figure 2. Combination of treatment groups in each animal was chosen at random, but the surgeon was not blinded during the procedure due to fixation plate variation. During surgery, one animal was sacrificed under anesthesia, and three limbs were left untreated due to plate complications for a final sample number of n=13, 11, 11 for stiff, early, and delayed, respectively. In all three groups, each defect received three high-density cell sheets contained within an electrospun, perforated, poly-(ε-caprolactone) (PCL) nanofiber mesh tube for a total of 6×10^6^ cells and 1.8 μg TGF-β1 per defect. In the group treated with BMP-2, the limbs were stabilized with a stiff fixation plate and 5 μg of recombinant human BMP-2 (rhBMP-2, R&D Systems) was delivered in an absorbable type I collagen sponge (DSM, Exton, PA), as previously described^32^. Animals were given subcutaneous injections of 0.04 mg/kg buprenorphine every 8 h for the first 48 h postoperative (post-op) and 0.013 mg/kg every 8 h for the following 24 h. All procedures were reviewed and approved by the Institutional Animal Care and Use Committee (IACUC) at the University of Notre Dame (Protocol No. 14-05-1778).

### Nanofiber mesh production

Nanofiber meshes were formed as described previously^23^ by dissolving 12% (w/v) poly-(ε-caprolactone) (PCL; Sigma-Aldrich, St. Louis, MO) in 90/10 hexafluoro-2-propanol/dimethylformamide (HFP/DMF) (Sigma-Aldrich, St. Louis, MO). The solution was electrospun at a rate of 0.75 ml/h onto a static aluminum collector. 9 mm × 20 mm sheets were cut from the product, perforated with a 1 mm biopsy punch (VWR), and glued into tubes around a 4.5 mm mandrel with UV glue (Dymax, Torrington, CT). Meshes were sterilized by 100% ethanol evaporation and washed 3x with sterile phosphate buffered saline (PBS; Lonza) before implantation.

### Fixation plate mechanical characterization

Characterization of the axial, torsional, and flexural plate stiffness was performed by screwing the plate onto two stainless steel 5/32” diameter rods through the tapped holes in the plate designed for bone screws. Axial tests were conducted in both the fixed-fixed and fixed-free configurations on the three plate configurations with a control rate of 0.02 mm/s to a displacement of 1.2 mm for the unlocked compliant and 0.005 mm/s to a displacement of 0.2 mm for stiff and locked compliant. Torsional tests were conducted in stiff, locked compliant, and unlocked compliant with a control rate of 0.1 deg/s to a displacement of 5 deg. Four-point bending tests were conducted using 5/32” square rods in the convex, concave, and side orientations on the three plate configurations with a control rate of 0.05 mm/s to a displacement of 2 mm for the unlocked compliant and 0.05 mm/s to a displacement of 1.5 mm for stiff and locked compliant. The stiffness under each loading condition was calculated as the slope of the linear region of the load-displacement curves.

### In vivo X-ray and microCT

*In vivo* X-rays and microCT scans were performed at 4, 8, and 12 weeks to determine bridging and assess bone volume of the defect respectively. *In vivo* CT scans were performed on an Albira PET/SPECT/CT system (Bruker, Billerica, MA) at 45 kVp, 0.4 mA, with best resolution (125 μm voxel size). 45 slices were analyzed in the center of each defect with a global threshold of 400 to determine bone volume. For the group treated with rhBMP-2 on an absorbable collagen scaffold, MicroCT analysis was performed using a Scanco μCT 80 system (Scanco Medical, Bassersdorf, Switzerland) at 70 kVp, 114 μA, at a resolution of 39 μm/voxel. For this group, 144 slices (144 slices × 39 μm/voxel = 5.616 mm) were analyzed in the center of each defect with a global threshold of 270 to determine bone volume. X-rays were taken using an Xtreme scanner (Bruker, Billerica, MA) at 45 kVp, 0.4 mA, with 2 second exposure time. A binary bridging was score was assigned by two independent, blinded observers, and determined as mineralized tissue fully traversing the defect.

### Ex vivo microCT

After 12 weeks the animals were euthanized by CO_2_ asphyxiation and hind limbs (n=10, 8, 8 for stiff, early, and delayed respectively after adjustment for limb complications and histology samples) were excised for high resolution microCT analysis using a Scanco μCT 80 system (Scanco Medical, Bassersdorf, Switzerland) at 70 kVp, 114 μA, at medium resolution (20 μm voxel size). 302 slices in the center of each defect were analyzed with a threshold corresponding to 488 mg HA/ccm, gauss filter width of 0.8, and filter support of 1.0. Two regions of interest (ROI) were analyzed; a 5 mm diameter circle centered on the medullary canal termed “defect”, and a “total” region in which any remaining bone outside the 5 mm defect was additively contoured. A third region, “ectopic”, was calculated by subtracting defect from total ROI. Bone volume, mineral density, bone volume fraction, polar moment of inertia (pMOI), and the morphometric parameters connectivity density, trabecular thickness, trabecular spacing, and trabecular number were calculated using a built-in algorithm that fits a maximal sphere in either the pore or trabecular strut at each voxel in the 3D space (Scanco evaluation script “V6”). Trabecular morphometry of three age-matched femoral heads were analyzed in the same manner for comparison. Proximal and distal total bone volume were calculated by halving the slice number in each sample and separately segmenting each half for comparison. All postmortem representative images were chosen based on average value for each illustrated parameter.

### Biomechanical testing

Femora (n=8, 7, 7 for stiff, early, and delayed respectively) excised at 12 weeks were biomechanically tested in torsion to failure. Limbs were cleaned of soft tissue and the fixation plate was carefully removed. Bone ends were potted in Wood’s metal (Alfa Aesar), mounted on a Bose ElectroForce biaxial load frame system (ELF 330, Bose EnduraTEC) and tested to failure at a rate of 3 degrees per second. For each sample maximum torque at failure was recorded and torsional stiffness was determined as the slope of a 5 degree linear region in the torque-rotation curve. Samples were compared to 7 age matched, un-operated femurs.

### Fibrin gel preparation and dynamic compression

At 80% confluency hMSCs were trypsinized and resuspended (P4) in a 10,000KIU/mL aprotinin solution (Nordic Pharma) with 19 mg/mL sodium chloride and 100mg/mL bovine fibrinogen (Sigma-Aldrich). This was combined 1:1 with a solution of 5U/mL thrombin and 40mM CaCl_2_ for a final solution of 50mg/mL fibrinogen, 2.5U/mL thrombin, 5000KIU/mL aprotinin, 17mg/mL sodium chloride, 20mM CaCl_2_, and 15×10^6^ cells/mL. Gel solution was pipetted into 5mm diameter × 2mm thickness cylindrical agarose molds to create uniform constructs with a total cell volume of approximately 600,000 each. Throughout the study, culture was maintained in chondrogenic media consisting of hgDMEM supplemented with 1% Penstrep, 100 KIU/mL aprotinin, 100 μg/mL sodium pyruvate, 40 μg/mL l-proline, 1.5 mg/mL bovine serum albumin, 4.7 μg/mL linoleic acid, 1x insulin-transferrin-selenium, 50 μg/mL L-ascorbic acid-2-phosphate, 100 nM dexamethasone (all Sigma-Aldrich), and 10ng/mL TGFβ3 (ProSpec-Tany TechnoGene Ltd., Israel). Fresh media was supplied every 3 days and culture was maintained in a humidified environment at 37°C, 5% CO_2_, and 5% O_2_. Dynamic unconfined compressive loading was applied to the constructs using a custom-made bioreactor. Load was applied 2 hours per day, 5 days a week at 1Hz and 10% strain after a .01N preload was applied. Load was applied either continuously for 5 weeks (early), for 2 weeks following a 3 week free swelling period (delayed), or for 2 weeks prior to a 3 week free swelling period (reversed), in comparison to 5 week free swelling controls.

### Biochemical analysis

Fibrin hydrogels were analyzed at 5 weeks for biochemical content (n=5). Samples were digested overnight at 60°C in a solution of 125 μg/mL papain, 0.1M sodium acetate, 5mM l-cysteine, 0.05M EDTA (all Sigma-Aldrich). DNA content was quantified with Hoechst Bisbenzimide 33258 dye assay (Sigma-Aldrich) as described previously,^58^ and sulfated glycosaminoglycan content was quantified using the dimethylmethylene blue dye-binding assay (Blyscan; Biocolor Ltd.). Orthohydroxyproline was measured by dimethylaminobenzaldehyde and chloramine T assay. Hydroxyproline to collagen ration of 1:7.69 was used to represent total collagen content.^59^

### Quantitative reverse transcription-polymerase chain reaction (qRT-PCR) analysis

hMSC sheet halves (n=3/group) and fibrin hydrogels (n=4-5/group) were homogenized in TRI Reagent (Sigma-Aldrich) for subsequent total RNA extraction and cDNA synthesis (PrimeScript^™^ 1st strand cDNA Synthesis Kit; Takara Bio Inc., Kusatsu, Shiga, Japan). One hundred nanograms of cDNA were amplified in duplicates in each 40-cycle reaction using a Mastercycler (Eppendorf, Hauppauge, NY) with annealing temperature set at 60°C, SYBR^®^ Premix Ex Taq^™^ II (Takara), and custom-designed qRT-PCR primers (Table I; Life Technologies, Grand Island, NY). Transcript levels were normalized to GAPDH and gene expression was calculated as fold change using the comparative C_T_ method.^60^

### Immunoblotting

hMSC condensations (n=3/group) were homogenized in CelLytic^™^ MT lysis buffer (Sigma-Aldrich) supplemented with Halt^™^ protease and phosphatase inhibitor cocktail (Thermo Scientific). Equal amounts (15 μg) of protein lysates, determined by standard BCA protein assay kit (Pierce; Thermo Fisher Scientific), were subjected to SDS-PAGE using 10% NuPAGE^®^ Bis-Tris gels (Invitrogen; Thermo Fisher Scientific) and transferred to 0.45 μm PVDF membranes (Millipore, Billerica, MA). Membranes were blocked with 5% bovine serum albumin in standard TBST. The phosphorylation of intracellular SMAD3 was detected using an anti-phospho-SMAD3 primary antibody (ab52903; Abcam, Cambridge, MA) followed by an HRP-conjugated secondary antibody (Jackson ImmunoResearch, West Grove, PA). Subsequently, the blot was stripped (Western Blot Stripping Buffer, Pierce; Thermo Fisher Scientific) and re-probed for the detection of total SMAD3 using an anti-SMAD3 primary antibody (ab40854; Abcam) and loading control using an anti-β-Actin primary antibody (A1978; Sigma-Aldrich) followed by their respective HRP-conjugated secondary antibodies (Jackson ImmunoResearch). Bound antibodies were visualized with the ECL detection system (Pierce; Thermo Fisher Scientific) on autoradiography film (Thermo Fisher Scientific). The intensity of immunoreactive bands was quantified using ImageJ software (NIH).

**TABLE I:**
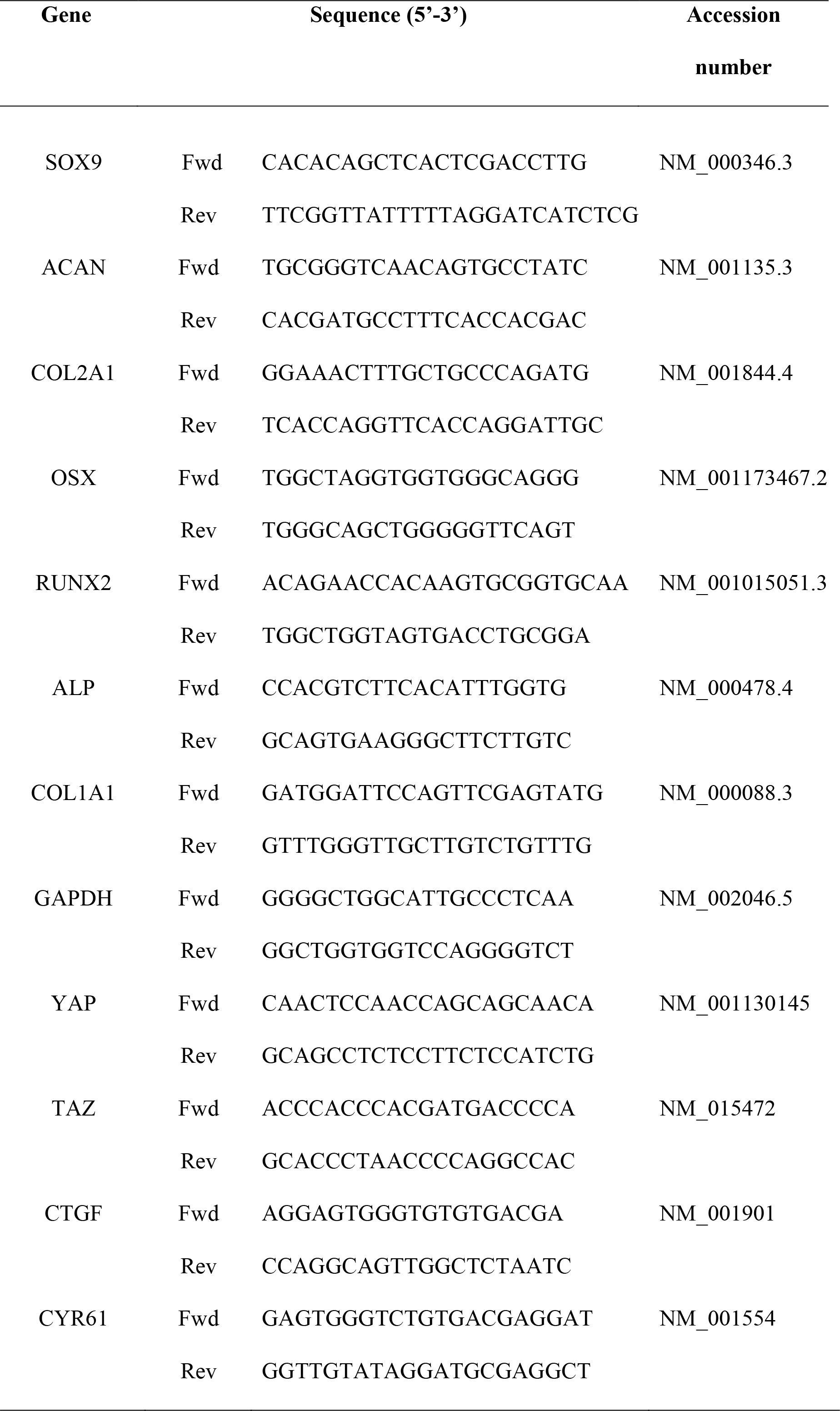
*Oligonucleotide primer sequences for qRT-PCR*.

### Histological analysis

hMSC sheet halves (n=3/group) were fixed in 10% neutral buffered formalin (NBF) for 24 h at 4°C before switching to 70% ethanol. Representative samples from each group were taken for histology at weeks 4 (n=4/group), 7 (n=3/group), and 12 (n=1/group) post-surgery, chosen based on microCT-calculated average bone or vascular volumes. Samples were fixed in 10% NBF for 72 h at 4°C and then transferred to 0.25 M ethylenediaminetetraacetic acid (EDTA) pH 7.4 for 14 d at 4°C under mild agitation on a rocker plate, with changes of the decalcification solution every 3-4 days. Following paraffin processing, 5-μm mid-saggital sections were cut using a microtome (Leica Microsystems Inc., Buffalo Grove, IL, USA) and stained with hematoxylin & eosin (H&E), Safranin-O/Fast-green (Saf-O), and picrosirius red (PSR; Polysciences, Inc., Warrington, PA). Light microscopy images, employing linearly polarized light for PSR-stained sections, were captured using an Olympus BX61VS microscope (Olympus, Center Valley, PA, USA) with a Pike F-505 camera (Allied Vision Technologies, Stadtroda, Germany).

Fibrin hydrogels were fixed in 4% paraformaldehyde (n=2) overnight at 4°C. Constructs were halved and paraffin embedded cut surface down, and sectioned at 5μm to provide a cross-section of the hydrogel center. Sections were cleared in xylene, rehydrated in graded alcohols, and stained for sGAG in 1% Alcian blue with 0.1% Nuclear Fast Red counter stain.

### Immunohistochemistry

Paraffin embedded hMSC sheets, fibrin hydrogels, and decalcified femurs were deparaffinized and rehydrated in successive incubations in xylene, ethanol, and diH2O. Heat induced antigen retrieval was performed for 8 minutes at 90°C in a Tris-EDTA antigen retrieval buffer; 10 mM Tris-base, 1 mM EDTA, 0.05% tween-20, pH 9.0. Endogenous peroxidase activity was quenched in .3% hydrogen peroxide in methanol. Non-specific binding was blocked using either serum-free protein block (DAKO; Santa Clara, CA, USA), for rabbit primary antibodies, or 2.5% horse serum from the RTU Vectastain Kit (Vector; Burlingame, CA, USA), for mouse primary antibodies. All primary antibodies were incubated overnight at 4°C. Rabbit primary antibodies were detected using Signal Stain Boost IHC Detection Reagent (Cell Signaling; Danvers, MA, USA) and mouse primary antibodies were detected using biotinylated universal secondary antibodies followed by incubation with streptavidin conjugated peroxidase from the RTU Vectastain Kit. Colorimetric antibody detection was performed using ImmPACT DAB peroxidase substrate kit (Vector) and counterstained with hematoxylin. Rabbit primary antibodies were yes-associated protein (YAP; D8H1X; Cell Signaling), transcriptional coactivator with PDZ-binding motif (TAZ; V386; Cell Signaling), collagen VI (col6a1; ab182744; Abcam) and rabbit isotype control IgG (IgG; DA1E; Cell Signaling). Mouse primaries were connective tissue growth factor (CTGF; ab6992; abcam; Cambridge, MA, USA), cysteine rich angiogenic inducer (CYR61; ab80112; abcam), collage type II (col2a1; sc-52658, Santa Cruz), and human nuclear antigen (HUNU; MAB1281; EMD Millipore; Darmstadt, Germany).

### Devitalized cell delivery

hMSC sheets (identical to previously described, using the same cells and growth factor presentation) were devitalized by 3 freeze/thaw cycles. Three devitalized sheets were placed into each electrospun, perforated, nano-mesh cylinder and placed in a critically-sized segmental defects (8mm) under stiff fixation (n=5). Limbs were assessed by *in vivo* microCT at 4, 8, and 12 weeks, and *ex vivo* microCT at 12 weeks for comparison with live cell delivery under stiff fixation.

### MicroCT angiography

Critically sized (8mm) segmental defect surgery was repeated in twenty 14 week-old male Rowett nude (RNU) rats (Charles River Labs, Wilmington, MA) under isoflurane anesthesia. In each rat a loaded limb, either early (n=10) or delayed (n=8) was paired with a contralateral control limb, stabilized by a stiff fixation plate that limited load transfer (stiff, n=18 total). Two animals were lost in the delayed group due to plate complications. Contrast enhanced microCT angiography was performed at week 3 in the early loading group and week 7 in the delayed group. Radiopaque contrast agent-enhanced microCT angiography was performed using a protocol modified from that described previously^48,61^ Briefly, the vasculature was perfused through the ascending aorta with sequential solutions of heparin, saline, 10% formalin, and lead chromate-based radiopaque contrast agent (Microfil MV-122, Flow Tech). After perfusion, limbs were excised and scanned via microCT (as described above) with both bone and contrast agent intact. Limbs were then decalcified with Cal-Ex II fixative/decalcifier (Fisher) for 2 weeks, scans were repeated, and subtraction was used to distinguish between bone and vessel parameters in a 5 mm diameter defect region of interest (ROI) and 7 mm diameter total ROI. Three representative samples from each group, chosen based on average microCT-computed vessel volume, were processed for histology.

### Cationic cartilage contrast-enhanced microCT

The cationic iodinated contrast agent, CA^4+^, was prepared as described previuosly,^62^ at a concentration of 24 mg I/mL, pH 7.4, and osmolality of 400 mOsm/kg. The paired early/stiff and delayed/stiff limbs, previously analysed by microCT angiography, were scanned before incubation and after 4 and 10 days of incubation. Day 10 scans were used for quantitative analysis. Limbs were incubated in 1.6 mL of CA^4+^) solution at room temperature. For the first hour of incubation and the last hour before each scan, sample tubes were incubated in a benchtop ultrasonic cleaner (model 97043-960, VWR) using a protocol modified from that previously described^50^. One stiff and one delayed limb (not paired) were lost from the delayed/stiff group due to tissue preparation error. MicroCT analysis was performed using a Scanco μCT 80 system (Scanco Medical, Bassersdorf, Switzerland) at 70 kVp, 114 μA, at a resolution of 15.6 μm/voxel. A region in the center of each defect was analyzed corresponding to the smallest defect in each group, 410 slices in early/stiff and 455 slices in stiff/delayed. Subtraction was used to distinguish between cartilage and vessel parameters in a 5 mm diameter “annulus” region of interest (ROI) and 1.5 mm diameter “core” ROI, centered on the medullary canal as determined by surgical screw placement in each sample. Registration of scans before and after incubation was performed using Analyze 10.0.

### Statistics

Bridging rates were assessed by chi-square test for trend (*p<0.05) and are presented as percent of samples bridged in each group; comparisons between groups were assessed with individual chi-squared tests and Bonferroni correction for multiple comparisons. Comparisons between groups were assessed by one- or two-way analysis of variance (ANOVA) with Tukey’s multiple comparison tests (*p<0.05), as appropriate. Raw data are displayed with mean ± s.d, mean ± s.e.m. or as box plots showing 25^th^ and 75^th^ percentiles, with whiskers at minimum and maximum values where specified. Where necessary and appropriate, data were log transformed to ensure normality and homoscedasticity prior to ANOVA. Normality of dependent variables and residuals were verified by D’Agostino-Pearson omnibus and Brown-Forsythe tests, respectively. For mechanical property regressions, we used an exhaustive best-subsets algorithm to determine the best predictors of maximum torque and stiffness from a subset of morphologic parameters measured, which included minimum or average polar moment of inertia (J_min_ or J_avg_), bone volume, bridging (binary score), and mineral density based on Akaike’s information criterion (AIC)^41^. The lowest AIC selects the best model while giving preference to models with less parameters. Finally, the overall best model for each predicted mechanical property was compared to its measured value using type II general linear regression.

In the devitalized cell experiment, surgical implantation and evaluation for this group was performed at a separate time from the live cell analysis and was therefore not included in quantitative statistical comparisons, though the presence of this group was accounted for in the ANOVA and post-hoc multiple comparisons shown in Figure 2c.

Differences in angiography between loaded samples and stiff samples were assessed by paired two-way student’s t-tests (*p<0.05), accounting for animal variability perfusion efficacy. Statistical comparisons were performed using GraphPad Prism (La Jolla, CA, USA). The sample sizes for microCT, mechanical testing, and contrast enhanced angiography analyses were determined with G*Power software^63^ based on a power analysis using population standard deviations and estimated effect sizes from our prior studies.^6,64^ The power analysis assumed a two-tailed alpha of 0.05, power of 0.8, and effect sizes of ranging from 0.1 to 0.3. A minimum sample number of n = 6 per group was computed, with an ideal sample number of n = 12 for all non-destructive and destructive analyses per time point. An n = 10 was selected for all non-destructive and destructive analyses per time point, accommodating a 5-10% complication rate consistent with our prior studies.

